# Natural Shortwave UV-B Radiation Increases UV absorbing pigments levels in Texas native grasses *Chasmanthium latifolium* and *Bouteloua curtipendula*

**DOI:** 10.1101/2022.12.30.519826

**Authors:** AN Nebhut, MR Semro, CA Worley, JR Shinkle

## Abstract

Plants are routinely exposed to UV-B radiation (280–315 nm) as a natural component of incident solar radiation. UV-B radiation is relevant to plants as both a source of damage and as a photomorphogenic cue, but the relative impacts of different wavelengths within the UV-B waveband are largely unresolved. Previous studies indicate that the full spectrum of solar UV-B radiation elicits unique responses in model plants under laboratory conditions compared to photons within the longwave UV-B band (300–315 nm), but the impacts of shortwave UV-B radiation (SW-UVB; 280–300 nm) remain unknown in a field environment. In this study, we examine the impact of shortwave UV-B radiation on the pigmentation and morphology of C_3_ shade-adapted inland sea oats (*Chasmanthium latifolium*) and C_4_ sun-adapted side-oats grama (*Bouteloua curtipendula*) in field conditions. We characterized the responses of these grasses to SW-UVB radiation by placing UV-naive individuals under SW-UVB excluding or transmitting filters at three field sites in Central Texas and measuring their responses through leaf pigment extract absorbance, leaf surface reflectance, and whole leaf chlorophyll and flavonoid content. We found that both species exhibited changes to their UV-B associated traits following exposure to the field environment, which were influenced by field site location, maximum daily temperature, and SW-UVB exposure. In particular, *C. latifolium* increased the ratio of its maximum to minimum absorbance in response to SW-UVB, while *B. curtipendula* exhibited higher flavonoid content in response to SW-UVB exposure. We believe this is the first time that the effects of SW-UVB radiation have been documented in a field environment.

## Introduction

### Plant responses to UV-B Radiation

Plants are routinely exposed to incident photons of ultraviolet-B radiation (UV-B) (280-315 nm) while seeking light for photosynthesis. While these photons compose only 0.5% of the solar radiation to reach the biosphere (Blumthaler 1993), UV-B radiation has an outsized effect on plants as a source of damage (Jansen et al. 1998; Jenkins 2009), as an environmental cue (Sampson and Cane 1999; Conner and Neumeier 2002; Strømme et al. 2015), and as a signal for morphological changes (Jenkins 2009).

One way plants sense the presence and fluence rates of UV-B radiation is through the photoreceptor UV RESPONSE LOCUS 8 (UVR8). UVR8 is found in diverse taxa, including green algae, bryophytes, lycophytes, and angiosperms, but the number of gene copies, consistency, and location of UVR8 expression varies between taxonomic groups (Robson et al. 2019; Tossi et al. 2019). While no additional UV-B photosensors have been definitively identified, there is evidence to suggest that plants may contain a second UV-photoreceptor in addition to UVR8. For example, *Arabidopsis* uvr8 mutants respond to UV-B radiation with the expression of UV-response genes and UV-B absorbing pigments (Tilbrook et al. 2013) and short and long wavelengths within the UV band elicit qualitatively different responses in various dicots, which indicates that distinct photosensory pathways are involved (Shinkle et al. 2004; Gardneret al. 2009).

UV-B exposure induces a number of responses in plants. One common and important UV-B associated response is the accumulation of UV-absorbing pigments in the vacuoles of epidermal cells, which act as a layer of molecular “sunscreen” by reducing the amount of UV-B radiation that reaches the UV-sensitive mesophyll cells, without attenuating the penetration of light in the visible spectrum (Li et al. 1993; Mazza et al. 2000; Caldwell et al. 2007; Bidel et al. 2007; Jenkins 2009). Some of these protective pigments, such as flavonoids, also serve as antioxidants and protect from oxidative damage and heat stress (Jansen et al. 1998; Searles et al. 2001; Caldwell et al. 2007; Bokszczanin et al. 2013). Another class of pigments known to respond to UV-B radiation is chlorophyll, which acts as both a general gauge of plant health (Smith et al. 2000) and as UV-B absorbing pigments (Bilger et al. 2001). However, rather than increasing with UV-B radiation, as would be expected from a protective pigment, plant chlorophyll content decreases under high UV-B conditions, which potentially contributes to plants’ reduced photosynthetic rate under enhanced UV-B radiation (Casati et al. 2002; Corria et al. 2005; Yao and Liu 2006). In addition to producing UV-B absorbing pigments, plants also protect sensitive tissues from UV-B radiation by reflecting UV-B photons from the leaf surface with structures such as thickened cuticular wax, trichomes, and altered cell optical properties (Jenkins 2009; Barnes et al. 2013; Robson et al. 2019).

Finally, there is considerable diversity in plant species’ responses to UV-B radiation. While some studies indicate that certain monocots are more resistant to UV-B stress than certain dicots (Pal et al. 2006; Kataria et al. 2013) and certain C_4_ grasses are more (Basiouny et al. 1978) or equally (Kataria et al. 2013) tolerant of UV-B radiation than certain C_3_ grasses, considerable interspecific diversity exists (Smith et al. 2000), possibly as a result of species niche, evolutionary history, and microclimate (Barnes et al. 2016). Despite this diversity, studies of plant responses to UV-B radiation are generally biased toward common model plants such as *Arabidopsis* and crop plants, rather than employing a wide range of species with diverse morphological, physiological, and life-history attributes. This bias may serve to exaggerate the effects of UV-B radiation because model and crop plants are often not adapted to the UV-B conditions in which they are placed (Krizek 2004; Robson et al. 2019).

Despite this depth of research on plant responses to UV-B radiation as a whole, little attention has been paid to the relative impact of different wavelengths within the UV-B waveband, including the impact of shortwave UV-B radiation (SW-UVB) (280–290 nm). Distinct responses to SW-UVB radiation may arise due to the high energy and thus relative danger of these photons compared to longwave UV-B (LW-UVB) (290–315 nm) photons, which may result in comparatively large impacts on plant growth and productivity (Flint and Caldwell 2003), or because the presence of SW-UVB fluctuates more throughout the season and the day than longer wavelengths (Kollias et al. 2003), making photons of these short wavelengths a useful environmental cue.

Already, unique responses to SW-UVB radiation have been shown in microarray gene expression studies in *Arabidopsis* (Ulm et al. 2004), growth analysis experiments in various dicots (Ros and Tevini 1995; Shinkle et al. 2004; Shinkle et al. 2005; Gardner et al. 2009), and UV-absorbing pigment absorption experiments on cucumbers and *Arabidopsis* (Shinkle et al. 2010), but no studies have thus far been conducted on the relevance of these wavelengths in the field environment with native plants.

### Importance of Studying Plant Responses to UV Radiation in a Natural Environment

Laboratory conditions often bear little resemblance to how plants experience UV-B radiation in a field environment. Greenhouse and laboratory studies frequently distort plant responses to UV-B radiation by limiting the spectral composition of the radiation to which plants are exposed, usually by conducting experiments under low photosynthetically active radiation (PAR), low UV-A, and high UV-B conditions (Krizek 2004; Robson et al. 2019). These unrealistic light environments remove plants’ ability to repair UV-B damaged DNA and weaken UV-B defenses, which leads to exaggerated UV-B damage and responses (Krizek 2004; Jenkins 2009; Robson et al. 2019). Likewise, whereas UV-B exposure in a laboratory setting is usually stable, UV-B radiation in a field environment fluctuates with a given location’s permanent attributes, such as latitude, altitude, and the surrounding landscape; with transient conditions such as the thickness of the ozone layer, meteorological conditions, surface reflectance, and the presence of atmospheric aerosols and pollutants; and with seasonal fluctuations in sunlight, temperature, and precipitation (Jansen et al. 1998; Jenkins 2009; Robson et al. 2019). Finally, solar UV-B radiation is only one of many environmental stressors that plants detect and respond to, and many plant responses to UV-B radiation also serve additional purposes in ameliorating other stressful conditions that are not present in laboratory studies. These other environmental variables may be long-term, such as rising temperatures over a summer (Coffey and Jansen 2019), or short-term, such as daily fluctuations in temperature (Csepregi et al. 2019). Together, overlap and cross-talk between plants’ responses to both UV-B radiation and other environmental variables cause plants to respond inconsistently to UV-B radiation across multiple years as average temperature, rainfall, etc. fluctuate (Callaghan et al. 2004; Robson et al. 2003; Hyyryläinen et al. 2015) and results in similar plants exhibiting different UV-B responses based upon their particular microclimatic circumstances (Teramura and Sullivan 1994).

The potentially exaggerated responses to UV-B radiation resulting from unrealistic laboratory conditions are of particular concern to those studying SW-UVB because while SW-UVB comprises a small fraction of the total solar radiation to reach the biosphere, it is also the portion of the solar spectrum most impacted by global ozone degradation (Kerr and McElrow 1993). This means that while SW-UVB radiation may have had little impact on plant UV-B associated responses in the past, its proportionally higher quantity compared to historical levels and the damage it produces relative to other wavebands means that SW-UVB radiation may be increasingly relevant to plant UV-B associated responses and overall physiology. Therefore, it is vital to accurately understand how plants respond to SW-UVB radiation in their natural light environments, not just to have a more complete picture of how plants sense and respond to their natural environment today, but also to project how the ecosystems of the future will respond to fluctuations in stratospheric ozone and climate change in the future.

### Aims and Hypotheses

Therefore, in this study, we conducted a series of SW-UVB exclusion experiments in the field to assess plant UV-B-associated responses as a result of (1) the presence of SW-UVB radiation and (2) maximum daily temperature. We tested these responses in two native Texas grass species chosen for their varying ecological niches: inland sea oats (*Chasmanthium latifolium*), a shade-adapted C_3_ species, and side-oats grama (*Bouteloua curtipendula*), a sun-adapted C_4_ species.

While we predicted that the plants exposed to SW-UVB radiation would produce a greater magnitude of UV-B associated responses than their unexposed counterparts, including increases in UV-B absorbing pigments, increases in leaf surface reflectance, decreases in chlorophyll content, and increases in flavonoid content, we expected that even SW-UVB unexposed plants in the field would produce some level of these UV-B associated responses as a result of overlap and cross-talk with other environmental variables. In particular, we expected that plants would adjust their UV-B associated responses to maximum daily temperature, with plants in higher-temperature environments exhibiting both greater responses to SW-UVB exposure (Csepregi et al. 2019) and higher UV-B associated responses overall. Finally, due to the differences in ecological niche and underlying physiological differences between *C. latifolium* and *B. curtipendula*, we expected that *C. latifolium*, as the C_3_ shade-tolerant species, would exhibit greater baseline UV-B pigmentation and morphology and produce greater UV-B associated responses following exposure to the field environment and SW-UVB than *B. curtipendula*, the C_4_ sun-adapted species.

## Methods

### Study Species and Care

We grew *B. curtipendula* and *C. latifolium* from commercial seed (Native American Seed, Junction, TX) in the UV-excluding Trinity University greenhouse. We planted the grasses in D40H Deepots (Stuewe & Sons, Inc., Tangent, OR) filled with a mixture of 50% potting mix (Garden-Ville, Austin, TX) and 50% Jiffy organic seed starting mix (Jiffy, Oslo, Norway). We watered the plants with deionized water as needed and allowed them to mature for 14 to 35 days after germination before moving them to our experimental set-ups in racks of up to 20 plants.

### Field UV-Exclusion Experiments

To test the relevance of SW-UVB radiation on plant UV-B associated responses, we conducted a series of five field experiments in the summer of 2019 at three privately-owned field sites: Bexar (latitude 29.462, longitude −98.484, and elevation 250 m), Kerr (latitude 30.038, longitude −99.391, and elevation 646 m), and Hays (latitude 30.141, longitude −97.874, and elevation 260 m). We selected these sites based on their availability through the Texas Ecolab Project, the availability of flat and open spaces, and ease of access. Following field site selection, we obtained the solar spectra of each location with an Apogee PS-200 spectroradiometer (Apogee Instruments, Logan, UT). In order to confine solar spectra readings to the UV region and eliminate spillover from visible wavelengths, we incorporated Hoya 330 UV bandpass filters into the sensors. Percent transmittance ranged from 90.7 to 91.7% from 290–320 nm. The Trinity University greenhouse transmitted no SW-UVB radiation, while each of the field sites experienced radiation as short as approximately 295 nm (*Fig. S3*).

At each field site, we constructed a single pair of UV-B exclusion cubes. Each pair was composed of two 1 m x 1 m x 1 m PVC pipe frames, one of which had an FS-UV transparent Aclar® (AC) filter (Electron Microscopy Sciences, Hatfield, PA) attached to the top and the other of which had an SW-UVB excluding 0.13 mm cellulose acetate (CA) filter (Piedmont Plastics, Rockville, MD) attached to the top. We placed the exclusion cubes approximately one meter apart from each other on a flat surface without overshading, tilted them southward along a north-south axis, reinforced them with baling wire, and then covered them in deer cloth to provide stability and protect against animal invasion.

After we constructed and reinforced the UV-exclusion cubes, we placed ten pots of *B. curtipendula* and *C. latifolium* under the UV-exclusion cubes and returned every four to five days to water the plants. On the fourteenth day, we took leaf samples, including ten leaves from each rack for pigment extracts, ten leaves from each rack for surface reflectance analysis, and 30 chlorophyll and flavonoid content surrogate readings from each rack, as described below. These samples were collected for each unique combination of plant rack, filter, treatment group, location, and species, and analyzed as described below. We repeated this experiment weekly for five consecutive weeks from June 4, 2019, to July 16, 2019, and collected daily maximum temperature data from Daymet (Thornton 2016).

### Leaf Sample Analysis

We chose the youngest fully expanded and visually healthy and intact leaves from each plug for each leaf sample. As grass leaves grow from the bottom up and UV-B associated responses vary along the length of a leaf due to differences in age, we took all samples from the middle of each leaf, under the assumption that these areas of the leaf were mature but not senescent. We analyzed these leaf samples through pigment extracts, reflectance spectra, and whole leaf chlorophyll and flavonoid content.

We produced leaf pigment extracts according to Shinkle et al. (2010). Ten leaves of each treatment group were first sectioned into 0.01-0.03 g segments and then placed in 1 mL of 99% methanol / 1% 12.1 M HCl in a dark freezer at −20°C. After 48 hours, we removed the leaf segments from the extract and allowed them to dry for 24 hours before weighing them. Immediately after removing the leaf segment, we measured the extract absorbance spectrum from 250 to 350 nm. We standardized these spectra by dry weight, then, to create a single quantification to represent the features of the absorbance spectra, we divided the maximum absorbance value by the minimum absorbance value to produce a maximum:minimum leaf extract absorbance ratio (MaxA:MinA) (see Mancinelli and Tolkowsky 1968, Tobin and Briggs 1969, and McArthur and Briggs 1970 for a similar technique used to track changes in phytochrome).

We obtained leaf reflectance spectra with a USB 2000+ reflectance spectrometer (Ocean Optics, Largo, FL), measured as a percentage reflection compared to a USRT-99-050-EPV white standard (Labsphere, North Sutton, NH). For each treatment group, we measured the upper surface of the middle of ten fresh leaves five times each, taking care to avoid the central vein. From the resulting spectra, we selected the leaf reflectance value at 300 nm as a representative value of the changes in leaf reflectance across the UV-B spectrum due to its central location and the linear shape of the leaf spectra as a whole.

We measured the whole leaf epidermal chlorophyll and flavonoids content through differential infrared transmission with a ForceA Dualex Scientific+™ meter (Dynamax Inc., Fresno, CA) whereby pigments absorbing in the UV-A range were determined by comparing chlorophyll fluorescence induced by a 375 nm excitation relative to a 640 nm reference excitation. We refer to the UV-A signal that represents flavonoids (“flavonoid surrogate”) as unitless FLAV values, as they represent a ratio of light excitation (Cerovic et al. 2012).

### Data Analysis

To compare baseline UV-B responses between species, we fit a set of linear mixed-effect models with each of the pretreatment UV-B variables as the outcome variable, a fixed effect of species, and a random effect of plant rack (*Equation 1*).

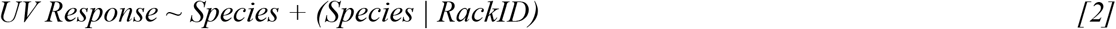

We then compared intraspecific UV-B responses resulting from exposure to the field environment (pre-vs. posttreatment measurements), filter (AC or CA), location (Bexar, Kerr, and Hayes), time (weekly groups from June 4, 2019, to July 16, 2019) and the average maximum daily temperature (°C) in the three days leading up to sample collection. To test these relationships, we fit a set of linear mixed-effect models with fixed effects of pre- or posttreatment measurement, the interaction between measurement and time, measurement and filter, measurement and temperature, and the three-way interaction between measurement, filter, and temperature, and random effects of rack ID and time (*Equation 2*).

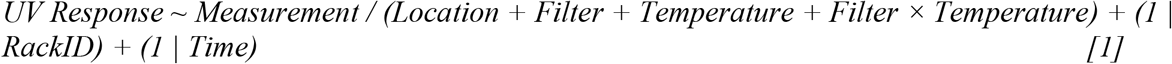

For each of these mixed-effect models, we started with the maximal model (i.e. the model with the most random effects-components), then progressively dropped terms as singularity or non-convergence occurred (Barr et al. 2013).

Prior to running the models, we confirmed that the model assumptions were met by visually checking for additivity and linearity between continuous independent and dependent variables. After selecting the maximal model for each UV response, we visually assessed whether the model met the homogeneity of variances and normality of residuals assumptions by visually checking the residual normal probability plot and the normal Q-Q plot, respectively. As the models met the assumptions, we proceeded without transformations. We assessed the whole model fit through piecewise structural equation modeling with R package piecewiseSEM (Lefcheck 2016) then analyzed the fixed effects with an ANOVA. For significant effects and interactions (alpha = 0.05), we used R package emmeans (Lenth 2020) to compare estimated marginal means and estimated marginal trends with a post hoc Tukey’s honest significance test difference test, utilizing p-value adjustments and Kenward-Rodger degrees of freedom.

To compare the relationship between MaxA:MinA and leaf surface reflectance before and after exposure to the field environment, we ran a Pearson correlation on the average MaxA:MinA and leaf surface reflectance values of *B. curtipendula* and *C. latifolium* by rack ID and measurement.

## Results

### Baseline UV-B Associated Responses

*B. curtipendula* and *C. latifolium* differ in their UV-B associated traits before exposure to the field environment (*Fig. 1*). *C. latifolium* maintains more UV-B absorbing pigments than *B. curtipendula*, as shown by its overall higher UV-absorbance spectra, and the two grasses contain different types of UV-absorbing pigments, as shown by the different shapes of their UV-absorbance spectra. While *B. curtipendula* has a peak at 320 nm and a trough at 285 nm, *C. latifolium* has a peak at 330 nm and a trough at 265 nm. *C. latifolium* has 85.2% higher MaxA:Min than *B. curtipendula* (*Fig. 1A*; F_1,29_ = 573.22; p < 0.001). In contrast, only the magnitude of the grasses’ leaf surface reflectance spectra is different, not the shape of the spectra, with *C. latifolium* exhibiting 26.5% higher leaf surface reflectance at 300 nm than *B. curtipendula* (*Fig. 1B*; F_1,29_ = 29.442, p < 0.001). Finally, *C. latifolium* contains 150.8% more chlorophyll (*Fig. 1C*; F_1,28.784_ = 141.68; p < 0.001) and 20.4% more flavonoids (*Fig. 1D*; F_1,28.224_ = 12.327; p < 0.001) than *B. curtipendula*.

**Figure 1.**
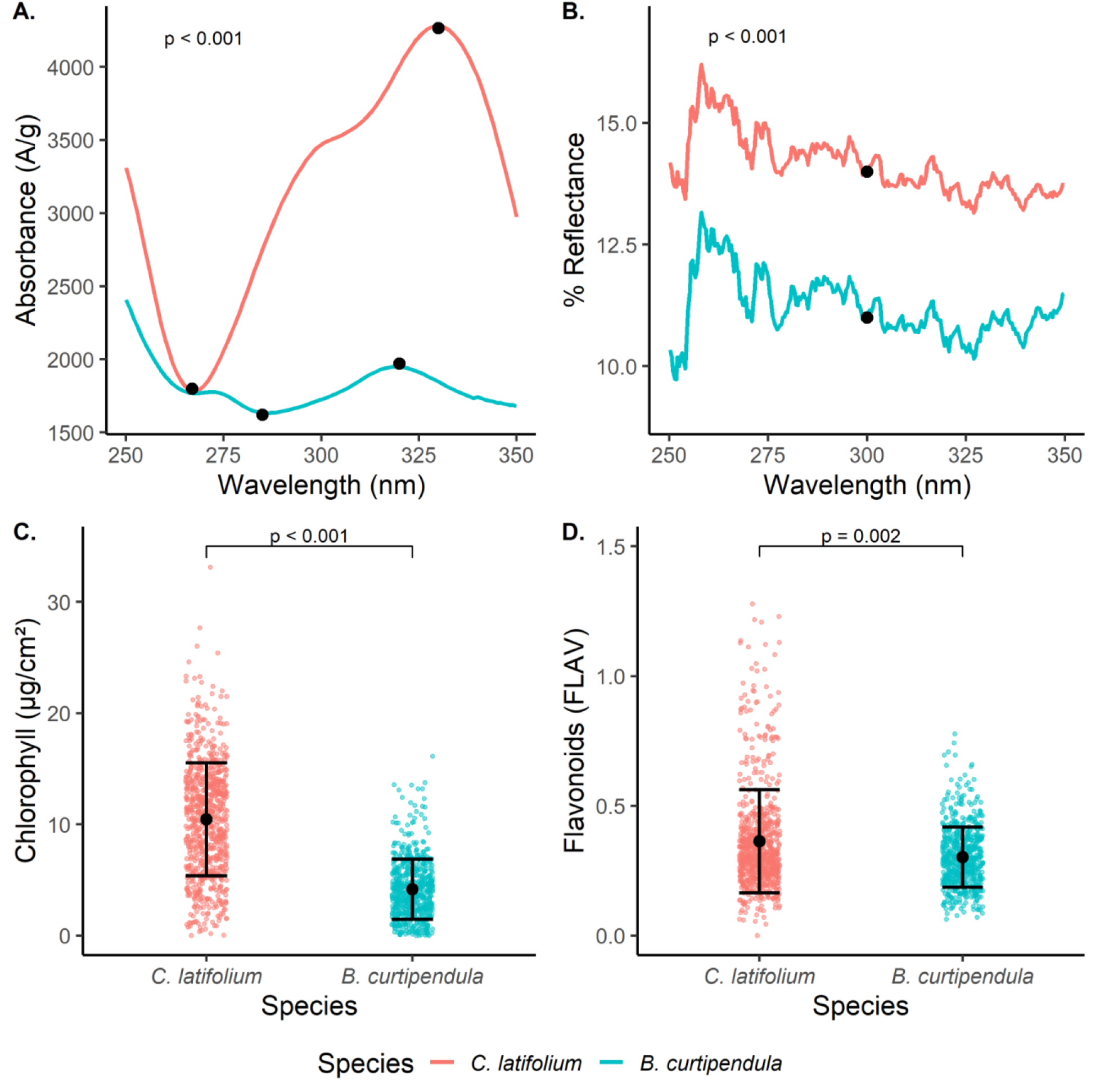
Baseline UV-B associated traits of *B. curtipendula* and *C. latifolium* prior to exposure to a field environment (*C. latifolium* in red, *B. curtipendula* in teal). A. Average pigment extract absorbance spectra of each species with highlighted maximum and minimum wavelengths; *B. curtipendula* has a peak at 320 nm and a trough at 285 nm and *C. latifolium* has a peak at 330 nm and a trough at 265 nm (n = 600). B. Average leaf surface reflectance spectra of each species with a highlighted “reference” wavelength; percent reflectance at 300 nm was chosen as the reference point for analysis (n = 3000). C. Chlorophyll content of whole leaf samples, shown are mean ± SD and data points (n = 1357). D. Flavonoid content surrogate of whole leaf samples, shown are mean ± SD and data points (n = 1381).

### UV-B associated traits respond to exposure to the field environment

Exposure to the field environment results in changes to the UV-B associated traits of both *B. curtipendula* and *C. latifolium*, but the character, direction, and magnitude of these changes differ by species (*Table 1*).

**Table 1.**
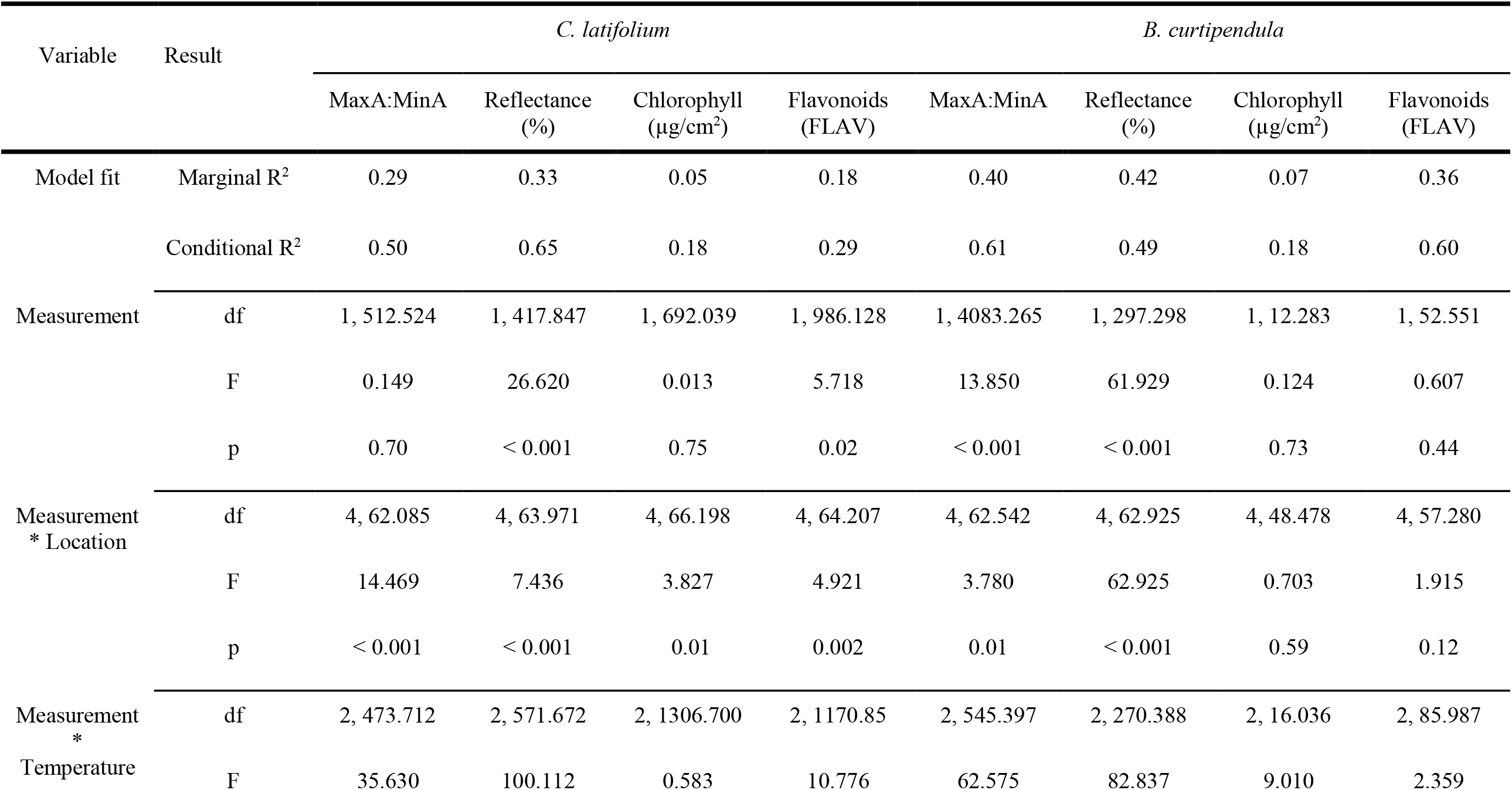
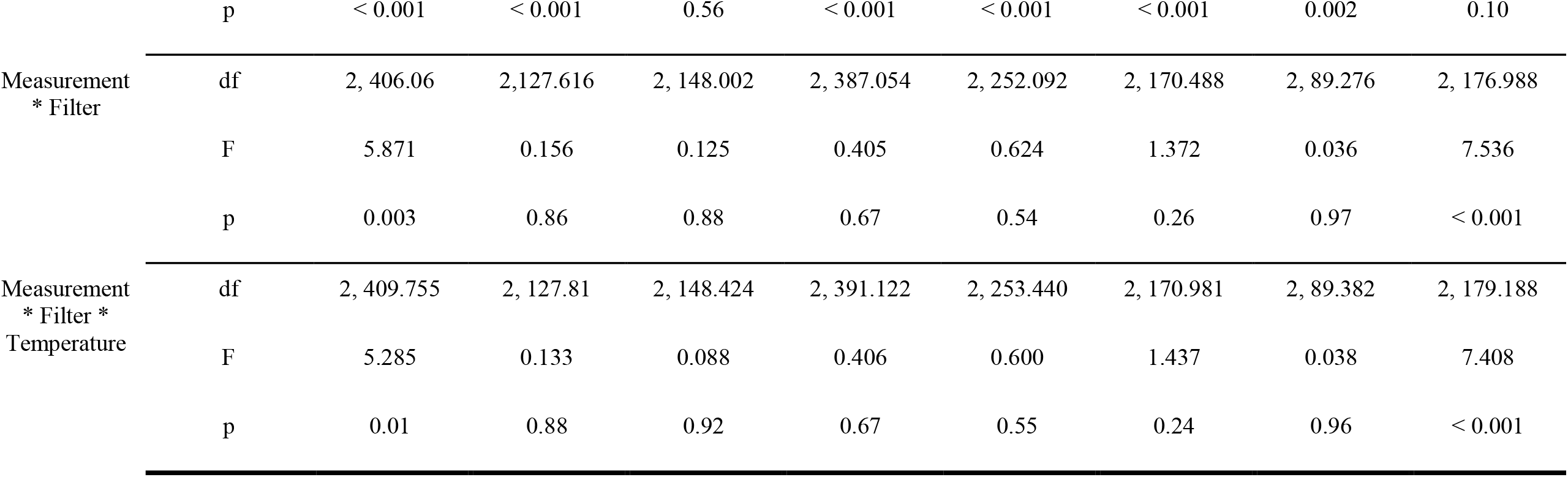
Model fit and ANOVA results of the linear mixed models of *C. latifolium* and *B. curtipendula* UV-B associated responses by measurement (pretreatment vs. posttreatment), the interactions between measurement and location (Bexar, Kerr, and Hays field sites), maximum daily temperature, or filter (AC or CA), and the three-way interaction between measurement, filter, and temperature.

*C. latifolium* responds to the field environment with a 28.5% decrease in leaf surface reflectance (F_1,418.85_ = 26.620; p < 0.001) and a 45.0% increase in flavonoid content surrogate (F_1,986.13_ = 5.718; p = 0.02), but no changes in MaxA:MinA (F_1,512.52_ = 0.149; p = 0.70) or chlorophyll content (F_1,692.04_ = 0.102; p = 0.75). *B. curtipendula* responds to the field environment with a 6.5% decrease in MaxA:MinA (F_1,483.27_ = 0.13.853; p < 0.001) and a 23.4% decrease in leaf surface reflectance (F_1,297.30_ = 61.929; p < 0.001), but no change in chlorophyll (F_1,12.28_ = 0.124; p = 0.73) or flavonoid content surrogate (*Fig. S1;* F_1,52.55_ = 0.607; p = 0.44). These differences are influenced by the field site location, with grasses at the different field sites exhibiting posttreatment UV-B associated responses of varying directionality and magnitude, independent of filter type or maximum daily temperature (*Fig. S2; Table S1*). Prior to exposure to the field environment, both *C. latifolium* and *B. curtipendula* exhibit a negative correlation between MaxA:MinA and leaf surface reflectance, but this correlation disappears after exposure to the field environment for both species (*Fig. 2*).

**Figure 2.**
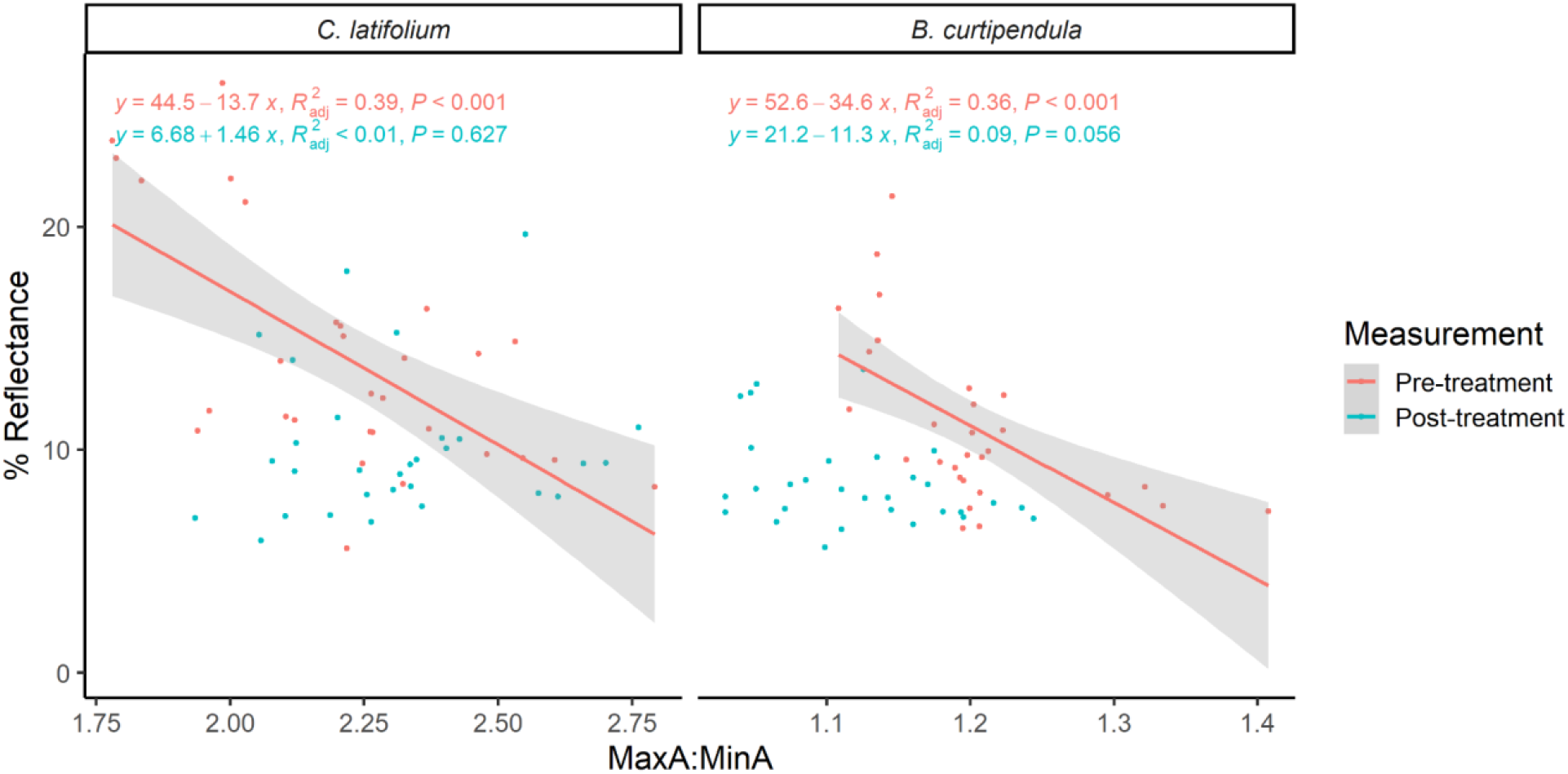
The relationship between treatment rack MaxA:MinA absorbance and leaf surface reflectance (%) in *C. latifolium* (n = 60) and *B. curtipendula* (n = 60) by treatment rack (pretreatment in red, posttreatment in blue). Shown are datapoints, line of best fit equation, Pearson’s correlation R^2^ and P-value, and, for significant correlations, the line of best fit and 95% confidence interval.

### Maximum daily temperature influences UV-B associated responses

The UV-B associated responses of *C. latifolium* and *B. curtipendula* are influenced by the average daily maximum temperature in the three days leading up to sample collection but are more sensitive to temperatures in the greenhouse environment than in the field.

*C. latifolium* exhibits significant interactions between temperature and MaxA:MinA, leaf surface reflectance, and flavonoid content surrogate, while *B. curtipendula* exhibits significant interactions between temperature and MaxA:MinA, leaf surface reflectance, and chlorophyll content (*Table 1*; *Fig. 3*). For both species, MaxA:MinA decreases with higher temperatures both in the greenhouse and after exposure to the field environment, but the pretreatment greenhouse response is of a larger magnitude (−0.31 ± 0.06 MaxA:MinA/°C for *C. latifolium* and −0.10 ± 0.01 MaxA:MinA/°C for *B. curtipendula*) than the posttreatment field response (−0.06 ± 0.01 MaxA:MinA/°C for *C. latifolium* and −0.01 ± 0.003 MaxA:MinA/°C for *B. curtipendula*). For leaf surface reflectance, both species exhibit a strong positive response to higher temperatures prior to exposure to the field environment (8.26 ± 0.92 %/°C for *C. latifolium* and 8.023 ± 0.70 8 %/°C for *B. curtipendula*), while after exposure to the field environment, both exhibit a smaller positive response (1.44 ± 0.18 %/°C for *C. latifolium* and 0.38 ± 0.14 %/°C for *B. curtipendula*). There is no significant interaction between measurement and temperature for *C. latifolium* chlorophyll content, while *B. curtipendula* chlorophyll content increases sharply with higher temperatures (1.11 ± 1.14 µg/cm^2^/°C) prior to exposure to the field environment and more shallowly (0.30 ± 0.22 µg/cm^2^/°C) after exposure to the field environment. Likewise, while there is no significant interaction between measurement and temperature for *B. curtipendula* flavonoid content surrogate, *C. latifolium* flavonoid content surrogate rises sharply with increasing temperature (0.10 ± 0.03 FLAV/°C) prior to exposure to the field environment and more shallowly (0.01 ± 0.006 FLAV/°C) after exposure to the field environment.

**Figure 3.**
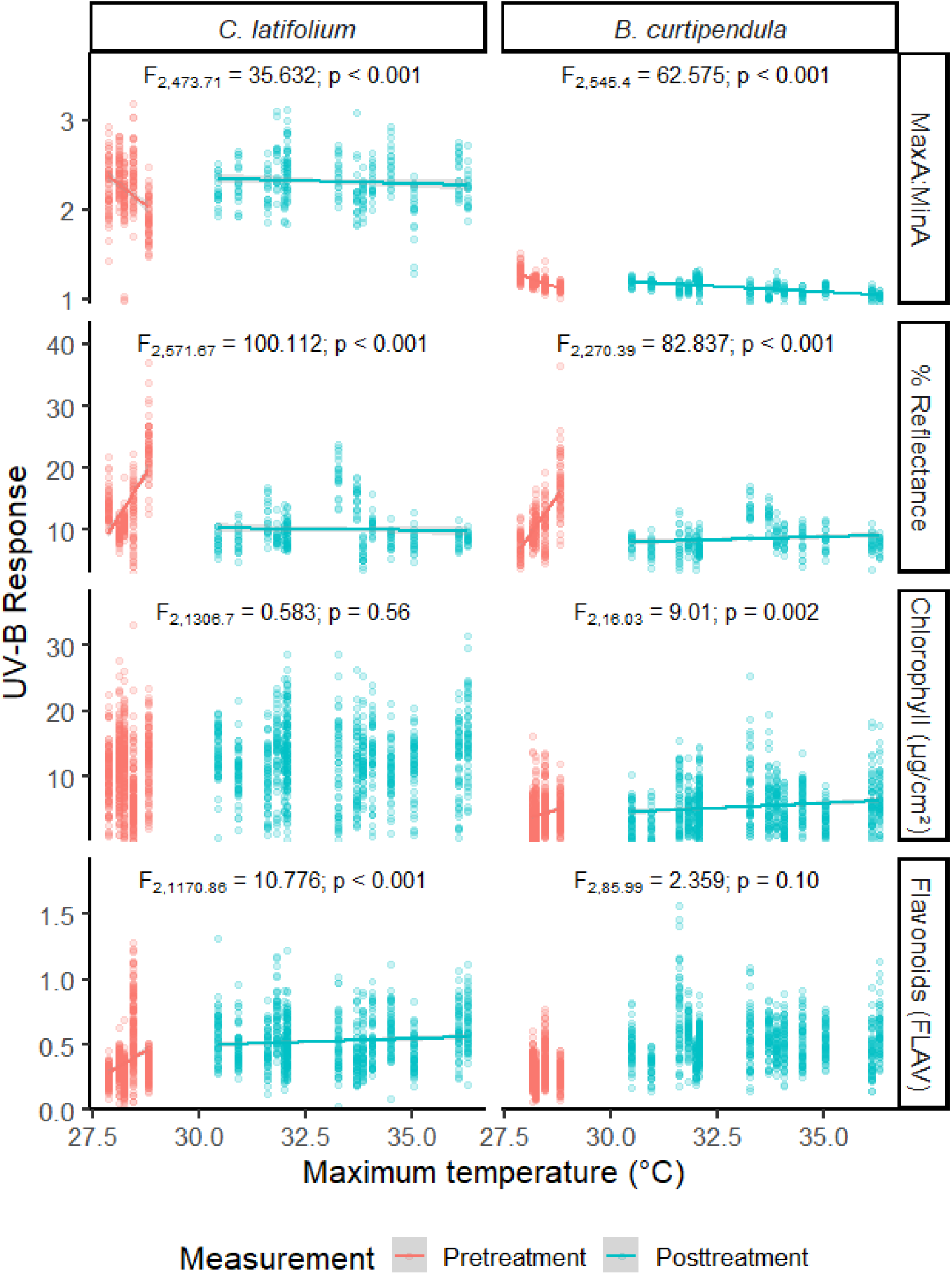
Relationship between three-day average maximum daily temperature (°C) and UV-B associated responses in *C. latifolium* (left) and *B. curtipendula* (right) before (red) and after (blue) exposure to the field environment. Shown are data points and ANOVA results and, for significant interactions, the lines of best fit and 95% confidence intervals.

### Exposure to natural SW-UVB radiation alters pigmentation

Exposure to natural SW-UVB radiation alters the MaxA:MinA of *C. latifolium* and the flavonoid content surrogate measurement of *B. curtipendula*, both alone and interactively with maximum daily temperature (*Table 1*).

For *C. latifolium*, there is a significant pretreatment difference in MaxA:MinA between the grasses assigned to AC and CA filters, with the AC-assigned grasses exhibiting 6.4% higher MaxA:MinA than the CA-assigned grasses. Posttreatment, the grasses placed under the UV-transparent AC filter exhibit a 6.9% increase in MaxA:MinA, whereas grasses placed under the UV-opaque CA filter exhibit only a 1.2% increase and have 12.3% lower MaxA:MinA than their UV-exposed counterparts (*Fig. 4A*; *Table 2*). Likewise, prior to exposure to the field environment, the grasses assigned to the CA filter respond to higher temperatures with larger decreases in MaxA:MinA (−0.47 ± 0.08 MaxA:MinA/°C) than the grasses assigned to the AC filter (−0.14 ± 0.08 MaxA:MinA/°C), but this difference disappears after exposure to the field environment, with the grasses assigned to the CA filter responding less negatively to higher temperatures (−0.04 ± 0.02 MaxA:MinA/°C), while the grasses assigned to the AC filter showed no significant change in their MaxA:MinA temperature response (−0.08 ± 0.02 MaxA:MinA/°C) (*Fig. 4B*; *Table 2*).

**Table 2.** Post-hoc Tukey’s honest significance difference test results for *C. latifolium* MaxA:MinA comparing estimated marginal means of the interaction between measurement (pretreatment and posttreatment) and filter (AC or CA) and estimated marginal trends of the three-way interaction between measurement, filter, and the three-day average maximum daily temperature (°C).

| Comparison | Result | Variable |  |
| --- | --- | --- | --- |
|  |  | Measurement *<br>Filter | Measurement *<br>Filter * Temperature |
| Pretreatment AC -<br>Posttreatment AC | df | 560.0 | 516 |
|  | t | -3.567 | -0.690 |
|  | p | 0.002 | 0.90 |
| Pretreatment CA -<br>Posttreatment CA | df | 560.0 | 516 |
|  | t | -6.372 | -4.826 |
|  | p | < 0.001 | < 0.001 |
| Pretreatment AC -<br>Pretreatment CA | df | 360.2 | 404 |
|  | t | 3.319 | 2.875 |
|  | p | 0.01 | 0.02 |
| Posttreatment AC-Posttreatment CA | df | 77.9 | 404 |
|  | t | 5.262 | -2.043 |
|  | p | < 0.001 | 0.17 |
| Pretreatment AC -<br>Posttreatment CA | df | 451.4 | 533 |
|  | t | -1.630 | -1.193 |
|  | p | 0.36 | 0.63 |
| Pretreatment CA -<br>Posttreatment AC | df | 451.4 | 533 |
|  | t | -7.802 | -4.485 |
|  | p | < 0.001 | 0.001 |

**Figure 4.**
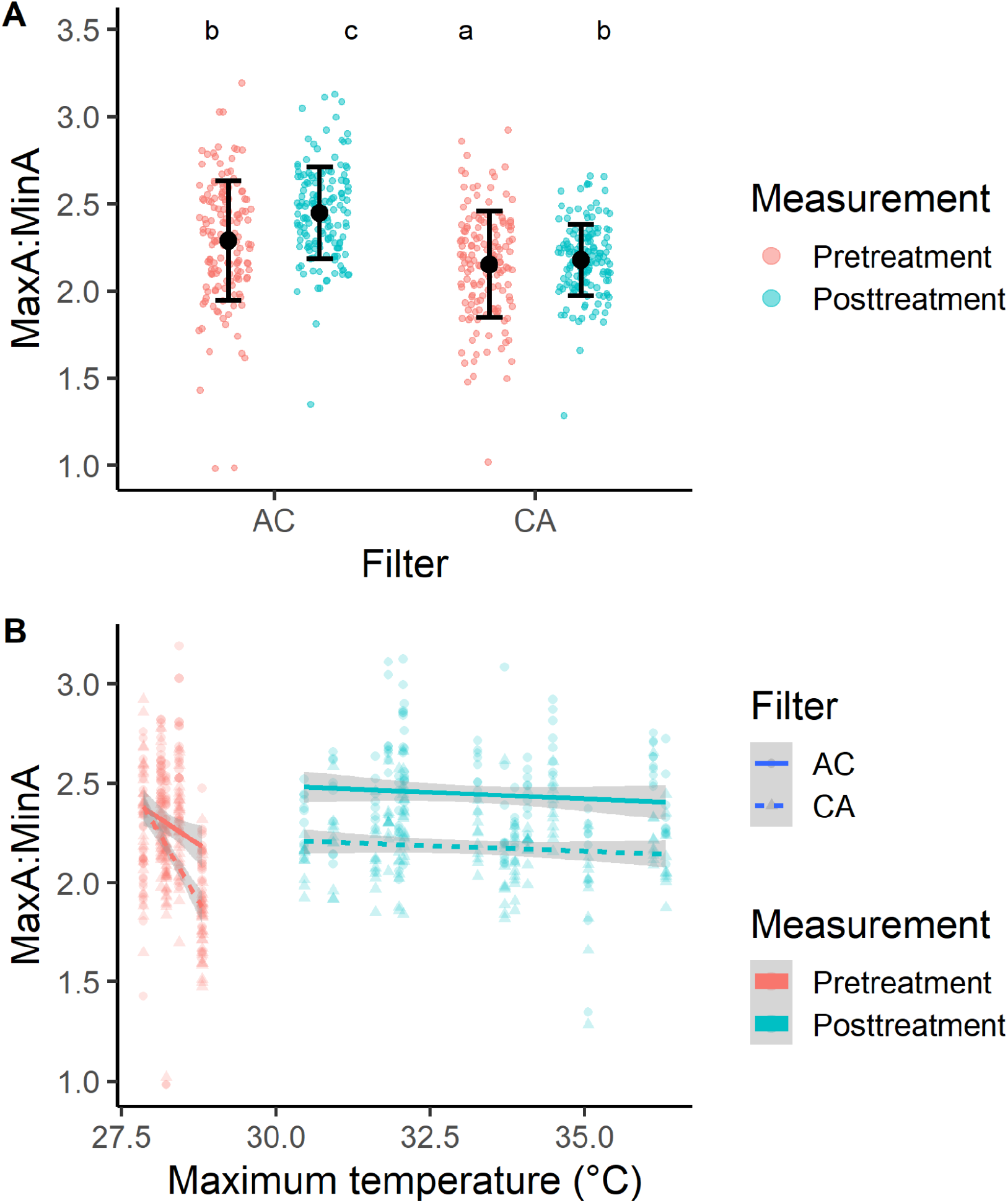
Impact of SW-UVB exposure on *C. latifolium* MaxA:MinA (n = 600). **A)** MaxA:MinA between plants under UV-transparent AC filters and UV-opaque CA filters before (red) and after (blue) exposure to the field environment. Shown are datapoints and mean ± SD and significance groups (treatments with the same letter code are not significantly different (Post-hoc Tukey’s honest significance test difference test of estimated marginal means, p > 0.05). **B)** Relationship between three-day average maximum daily temperature (°C) and MaxA:MinA between plants under UV-transparent AC filters (circles, solid lines) and UV-opaque CA filters (triangles, dashed lines) before (red) and after (blue) exposure to the field environment. Shown are data points, the lines of best fit, and 95% confidence intervals.

For *B. curtipendula*, the grasses placed under the UV-transparent AC filter and UV-opaque CA filter exhibit no significant differences in flavonoid content surrogate from each other prior to treatment and no significant difference in flavonoid content surrogate between pretreatment and posttreatment sampling, but posttreatment grasses under the UV-transparent AC filter exhibit 3.0% higher flavonoid content surrogate than posttreatment grasses under the UV-opaque CA filter (*Fig. 5A*; *Table 3*). Prior to treatment, *B. curtipendula* exhibits no differences in flavonoid responses to temperature between the grasses assigned to the two filter groups, and following treatment, there are no significant differences in their flavonoid temperature responses before and after treatment for either group. Following treatment, however, UV exposure altered the grasses’ responses to high temperatures, as the grasses under the UV-transparent AC filter decreased their flavonoid content surrogate in response to higher temperatures (−0.03 ± 0.02 FLAV/°C) while the grasses under the UV-opaque CA filter increased their flavonoid content surrogate in response to higher temperatures (0.01 ± 0.02 FLAV/°C) (*Fig 4B*; *Table 3*).

**Table 3.** Post-hoc Tukey’s honest significance test difference test results for *B. curtipendula* flavonoid content (FLAV) comparing estimated marginal means of the interaction between measurement (pretreatment and posttreatment) and filter (AC or CA) and estimated marginal trends of the three-way interaction between measurement, filter, and the three-day average maximum daily temperature (°C).

| Comparison | Result | Variable |  |
| --- | --- | --- | --- |
|  |  | Measurement *<br>Filter | Measurement *<br>Filter * Temperature |
| Pretreatment AC -<br>Posttreatment AC | df | 64.0 | 53.2 |
|  | t | -0.738 | 0.662 |
|  | p | 0.86 | 0.91 |
| Pretreatment CA -<br>Posttreatment CA | df | 64.5 | 53.5 |
|  | t | -1.318 | -0.470 |
|  | p | 0.56 | 0.97 |
| Pretreatment AC -<br>Pretreatment CA | df | 119.3 | 116.6 |
|  | t | 0.958 | 1.091 |
|  | p | 0.77 | 0.70 |
| Posttreatment AC -<br>Posttreatment CA | df | 37.7 | 84.7 |
|  | t | 2.697 | -3.325 |
|  | p | 0.05 | 0.01 |
| Pretreatment AC -<br>Posttreatment CA | df | 71.5 | 47.8 |
|  | t | -0.400 | 0.337 |
|  | p | 0.98 | 0.99 |
| Pretreatment CA -<br>Posttreatment AC | df | 72.2 | 48.0 |
|  | t | -1.630 | -0.140 |
|  | p | 0.37 | 1.00 |

**Figure 5.**
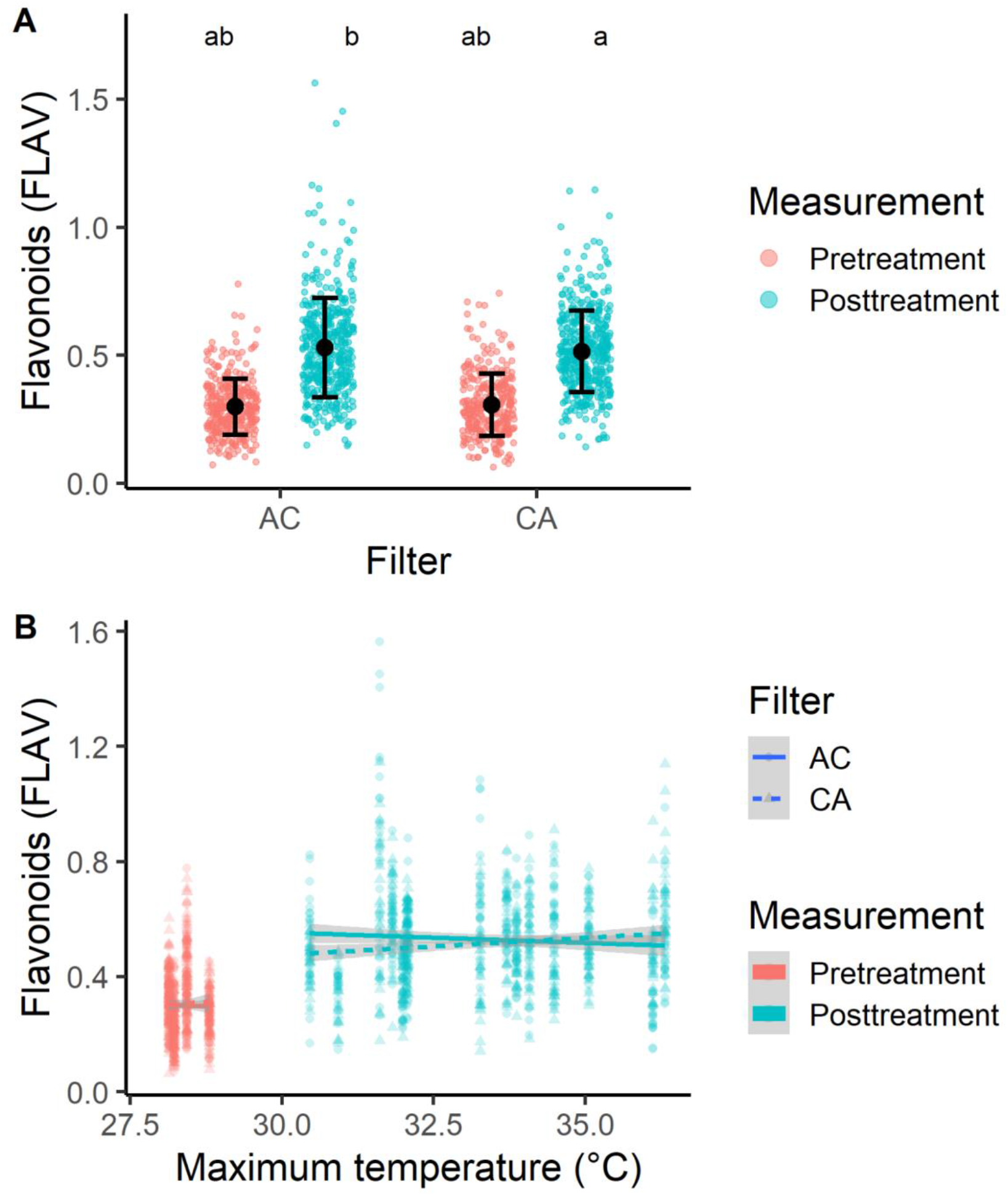
Impact of SW-UVB exposure on *B. curtipendula* flavonoid content surrogate (n = 1498). **A)** Flavonoid content surrogate between plants under UV-transparent AC filters and UV-opaque CA filters before (red) and after (blue) exposure to the field environment. Shown are datapoints and mean ± SD and significance groups (treatments with the same letter code are not significantly different (Post-hoc Tukey’s honest significance test difference test of estimated marginal means, p > 0.05). **B)** Relationship between three-day average maximum daily temperature (°C) and flavonoid content surrogate between plants under UV-transparent AC filters (circles, solid lines) and UV-opaque CA filters (triangles, dashed lines) before (red) and after (blue) exposure to the field environment. Shown are data points, the lines of best fit, and 95% confidence intervals.

## Discussion

When greenhouse-grown *B. curtipendula* and *C. latifolium* are placed in a field environment, both species exhibit changes in their UV-B protective pigmentation and morphology. Several environmental cues or stressors contribute to the plants’ UV-B associated responses, including temperature and the presence of SW-UVB radiation. We believe this is the first time that the effects of SW-UVB radiation have been documented in a field environment. To properly frame our findings, we acknowledge that the experimental design involved a transition where UVB naïve plants were abruptly moved into environments with full solar UVB. For that reason, our observations demonstrate that plants have the capacity to sense and respond to SW-UVB, but not how they do so when grown in natural light environments for their whole life cycle.

### UV-B associated traits exhibit unintuitive responses to exposure to the field environment

Upon exposure to a field environment and its stressors, we expected that the grasses would display typical UV-B associated responses in the form of increases in MaxA:MinA, leaf surface reflectance, and flavonoid content, paired with a decrease in chlorophyll content. Instead, while *C. latifolium* followed our predictions by raising its MaxA:MinA following exposure to a field environment, indicating an increase in UV-absorbing pigments, exposure to a field environment had the opposite effect on *B. curtipendula*, which dropped its MaxA:MinA, indicating a loss of pigmentation. Likewise, while leaf surface reflectance was expected to increase following exposure to a field environment as a defensive measure to prevent UV-B radiation from reaching the sensitive mesophyll cells, leaf surface reflectance actually decreased in both grass species.

Notably, both species show a negative relationship between MaxA:MinA and leaf surface reflectance prior to exposure to the field environment. This negative relationship may result from a potential trade-off between these two forms of UV-B defense, whereby individuals investing in energetically expensive UV-B absorbing pigments do not need to produce abundant leaf waxes or other UV-B reflective structures and vice versa. This trade-off between UV-absorbing compounds, including UV-absorbing photosynthetic compounds such as carotenoids and chlorophyll, and leaf reflectance is also found in Li et al’s (2019) assessment of UV-associated traits in four alpine broadleaf herbaceous species along an altitudinal gradient, which suggests that the trade-off is taxonomically widespread. However, its loss following exposure to the field environment suggests that in a stressful environment, these grasses may opt to allocate resources to both aspects of UV-B defense, rather than relying upon one or the other.

While we anticipated that both species would decrease their chlorophyll content following exposure to a field environment as a result of stress, exposure to the field environment had no measured impact on either species’ chlorophyll content. This finding diverges from results showing that plants lower their chlorophyll content as either a stress response following exposure to UV-B (Casati et al. 2002; Yao and Liu 2006) or as a trade-off for energetically expensive UV-absorbing compounds (Barnes et al. 2015). The stable chlorophyll content in both *C. latifolium* and *B. curtipendula* might indicate an acclimation response because chlorophyll is UV-B absorptive and therefore could be used to protect underlying tissues (Li et al. 2019).

Finally, while we expected that exposure to the field environment would increase flavonoid content in both species, this was only observed in *C. latifolium*, while *B. curtipendula* exhibited no change in flavonoid content surrogate.

Throughout the experiments, the UV-B associated responses of *B. curtipendula* and *C. latifolium* varied between treatment groups that differed in location of their placement. These variations could be a result of many factors, including variations in the surrounding landscape, meteorological conditions, and the presence of aerosols and pollutants along the urban-rural gradient captured by our field sites (Jansen et al. 1998; Jenkins 2009; Robson et al. 2019). It is not possible to determine which of these mechanisms is at play from our study design, but each of these variables warrant exploration in future studies.

### High temperatures alter UV-B associated traits

We anticipated that high daily maximum temperatures in the three days leading up to sample collection, as a source of environmental stress, would result in decreases in chlorophyll and increases in MaxA:MinA, leaf surface reflectance, and flavonoid content. Of these predictions, only the increase in leaf surface reflectance and the increase in *C. latifolium* flavonoids were supported by our data, whereas both species exhibited decreasing MaxA:MinA in response to higher temperatures, *B. curtipendula* increased its chlorophyll content in response to higher temperatures while *C. latifolium* chlorophyll content was not influenced by temperature, and *B. curtipendula* flavonoid content surrogate was not influenced by temperature. The observed decrease in MaxA:MinA in both species under high temperatures may be explained by heat stress and parallels findings that high temperatures decrease pigmentation in wine grapes (Mori et al. 2005; Mori et al. 2007; Rosas et al. 2017). Likewise, while high temperatures are typically expected to decrease chlorophyll content as a stress response, the increase in *B. curtipendula* chlorophyll content in response to higher temperatures may result from the grasses increasing chlorophyll production to make up for losses in photosynthetic rate per unit of chlorophyll at these high temperatures (Sayed et al. 1989; McDonald and Paulsen 1996).

While the UV-B associated responses of both *C. latifolium* and *B. curtipendula* were sensitive to maximum daily temperature in the three days leading up to sample collection both before and after exposure to the field environment, the grasses were more sensitive to high temperatures in the greenhouse environment than in the field environment, as shown by the more extreme relationships between UV-B associated responses and temperature in the pretreatment group compared to the posttreatment group. This result may be explained by the role of compounding stressors. While some stressors may interact synergistically to increase a plant’s response to both, such as the drought and UV-B stress synergistically increasing phenolics production (Caldwell et al. 2007), in other cases, a dampening effect occurs, whereby exposure to a stressor, such as falling winter temperatures, decreases a plant’s response to a second stressor, such as UV-B radiation (Coffey and Jansen 2019). Given the many potential stressors of the field environment that may be prompting the grasses to respond less to high temperatures in the field than in the greenhouse environment, such as insect herbivory (Izaguirre et al. 2007), water stress (Balakumar et al. 1993), and changes in the spectral composition and amount of solar insolation (Flint and Caldwell 2002), further research is required to elucidate how the various drivers of plant UV-B associated responses interact with each other in a field environment to produce a coherent response.

### Natural SW-UVB radiation increases pigmentation in *C. latifolium* and *B. curtipendula*

Baseline UV-B associated traits varied by species, with *C. latifolium* maintaining higher baseline UV-B defenses in the form of high absorbance values and reflectance values across the UV spectrum, and high chlorophyll and flavonoid content, compared to *B. curtipendula*, which maintained lower absorbance, reflectance, and chlorophyll and flavonoid content. This result was unexpected, given our prediction that *C. latifolium*, as a shade-tolerant species, would have lesser UV-B defenses compared to the sun-adapted *B. curtipendula*.

These species also responded differently to SW-UVB exposire, with *C. latifolium* exhibiting higher MaxA:MinA and no SW-UVB-prompted changes in temperature sensitivity, and *B. curtipendula* exhibiting overall higher flavonoid surrogate content and a slight negative relationship between flavonoid content surrogate and temperature. This is in contrast to the slight positive relationship between flavonoid content surrogate and temperature exhibited by their unexposed counterparts. These increases in pigmentation were expected given the protective qualities of flavonoids and other UV-B absorbing pigments and likely represent a defense mechanism by the grasses against the harms wrought by UV-B radiation (Li et al. 1993; Mazza et al. 2000; Caldwell et al. 2007; Bidel et al. 2007; Jenkins 2009). The negative relationship between *B. curtipendula* flavonoid content surrogate and temperature, following SW-UVB exposure, suggests that the compounding stressors of SW-UVB and high temperatures decreased pigmentation production.

These findings do not support our hypothesis that as a shade-adapted C_3_ plant, *C. latifolium* is more sensitive to UV-B radiation and would, therefore, exhibit greater changes in UV-B associated pigmentation and morphology than the sun-adapted C_4_ plant *B. curtipendula*. Simply put, instead of responding to UV-B radiation with changes in morphology and pigmentation that differ only their magnitude, the species exhibit qualitatively different responses involving changes in different suites of pigments, possibly suggesting diverging UV-B protective “strategies.” However, further research is required to explore which pigments underlie the species’ UV-B responses and the evolutionary mechanisms behind these differences.

Overall, these differences in pigment extract absorbances and leaf surface reflectances between plants that were and were not exposed to SW-UVB radiation highlight the ability of plants to fine-tune their UV-B responses to their particular environment and confirm that plants do sense and respond to SW-UVB radiation in a natural light environment.

### Applications

Our finding that the presence of SW-UVB radiation impacts the UV-B associated responses of both *B. curtipendula* and *C. latifolium* confirms earlier studies indicating that these high-energy photons have distinct impacts on plant UV-B associated responses and indicates that plants are sensitive to SW-UVB radiation under natural conditions (Ros and Tevini 1995; Ulm et al. 2004; Shinkle et al. 2004, 2005, 2010; Kalbina et al. 2008; Gardner et al. 2009). This result is of particular interest given the changes in the quantity of SW-UVB radiation reaching the biosphere as a result of stratospheric ozone depletion. While the ozone layer is recovering from its low point in the 1990s, full recovery remains many decades away due to the protracted lifespans of chlorofluorocarbons and halons (Chipperfield et al. 2017), meaning that plants will continue to encounter heightened SW-UVB exposure for decades and must adjust their UV-B associated responses accordingly.

A more complete understanding of plant UV-B associated responses within the context of their environment is crucial because UV-B associated responses impact not just the plants themselves, but also many of the competitive relationships between individuals and species that together form the basis of ecosystems as a whole (Barnes et al. 1996). Furthermore, as both UV-B radiation and the other factors that shape plant UV-B associated responses, such as temperature, are disrupted by climate change and atmospheric ozone depletion, the plants of the future will exhibit different UV-B associated responses than the plants of today. These differences will, in turn, ripple throughout ecosystems in manners that will often be difficult to predict. As a telling example, increased UV-B radiation in Northern Fennoscandia impacts the chemical composition of forage to decrease foraging quality (Martz et al. 2011). Other changes to plant UV-B associated responses and resulting ecosystem impacts driven by climate change and ozone depletion have already been cataloged by several reviews, including Ballaré et al. 2011, and are likely to become more severe as these processes progress.

## Supporting information

All supplements

## Acknowledgements

This research was supported by the Texas Ecolab Project and the Murchison Research Fellowship from Trinity University. We note our appreciation to Calvin Dao for writing our reflectance data compilation software and to Troy Murphy for the use of his reflectance spectrometer. Kendall Kotara, Sarah Pickett, and Emily Welp performed pilot experiments that informed this work.

## Abbreviations

(SW-UVB): shortwave UV-B radiation
(CA): SW-UVB excluding cellulose acetate filter
(AC): SW-UVB transparent aclar filter
(MaxA:MinA): maximum:minimum leaf extract absorbance ratio
(FLAV): flavonoid content surrogate unit

---

December 6, 2022

Dear Miquel Vall-Llosera Camps and PLOS editorial staff,

This letter accompanies the submission of the manuscript “Natural Shortwave UV-B Radiation Increases UV absorbing pigments levels in Texas native grasses *Chasmanthium latifolium* and *Bouteloua curtipendula*” by Nebhut, Semro, Worley, and Shinkle. To our knowledge, this is the first report of plant responses to short-wavelength UVB radiation (less than 300 nm) using solar radiation and exclusion filters under field conditions. It is also the first report of monocot responses to short wavelength UVB that we can find since 1956. Over the past 36 years, there have been multiple reports of plant responses specifically to short-wavelength UVB in laboratory conditions using artificial light sources. These studies used several dicot species under both etiolated and light-grown conditions. Given the integration of multiple signals and stressors by plants grown under field conditions, the previous studies leave open the possibility that plants do not exhibit unique responses to short-wavelength UVB outside of the laboratory. The current study goes some ways to answering that question while also documenting other environmental variables that contribute to the range of responses studied. Based on the foregoing description, this manuscript is submitted as a research article. It falls in the general area of plant physiological ecology and/or plant photobiology. Having considered publication of this work in other journals, I identified several potential reviewers. These are:

Stephan Flint

Rainer Hofmann

Marcel Jansen

Marta Rieriestè

T Matthew Robson

The history of the authors’ interactions with PLOS ONE is as follows: In 2019, after the ASPB meeting, PLOS ONE representative Jamie Males emailed me to encourage me to submit the findings from our poster once it was ready for publication. There were many reasons for the delay, but a major non-COVID reason was that we, in consultation with experts, made the statistical analysis more comprehensive. This did not change the results’ implications but gave us more confidence in them. After presenting the revised analysis at Plant Biology 2022, I emailed Dr. Males at PLOS ONE to restart the discussion about the manuscript. The first person to reply was Sebastian Shepherd, who was also encouraging about a manuscript submission. This November, I heard from Alejandra Clark that Dr. Shepherd had left PLOS ONE and that the most likely editor was Miquel Vall-Llosera Camps.

The only “opposed” reviewer might be Dr. Paul Barnes. He served on Ms. Nebhut’s undergraduate thesis committee and so might have a conflict of interests. I will leave that for PLOS ONE to decide.

Thank you for considering our work.

Sincerely,

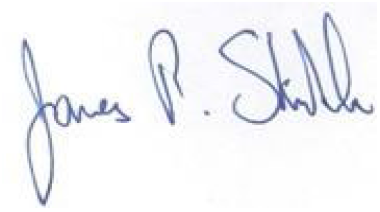

Dr. James Shinkle

Professor of Biology

Trinity University

## Supporting Information

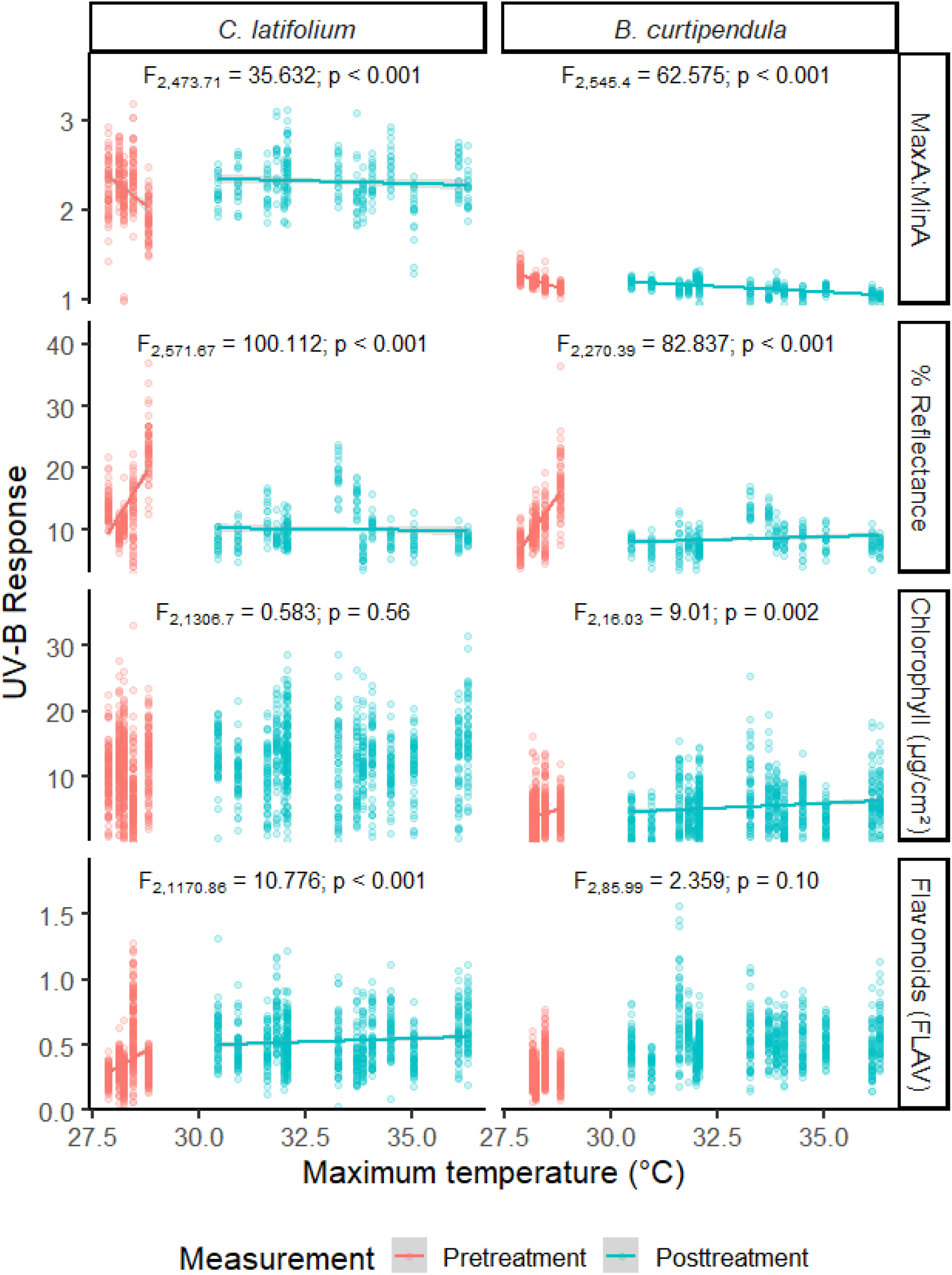

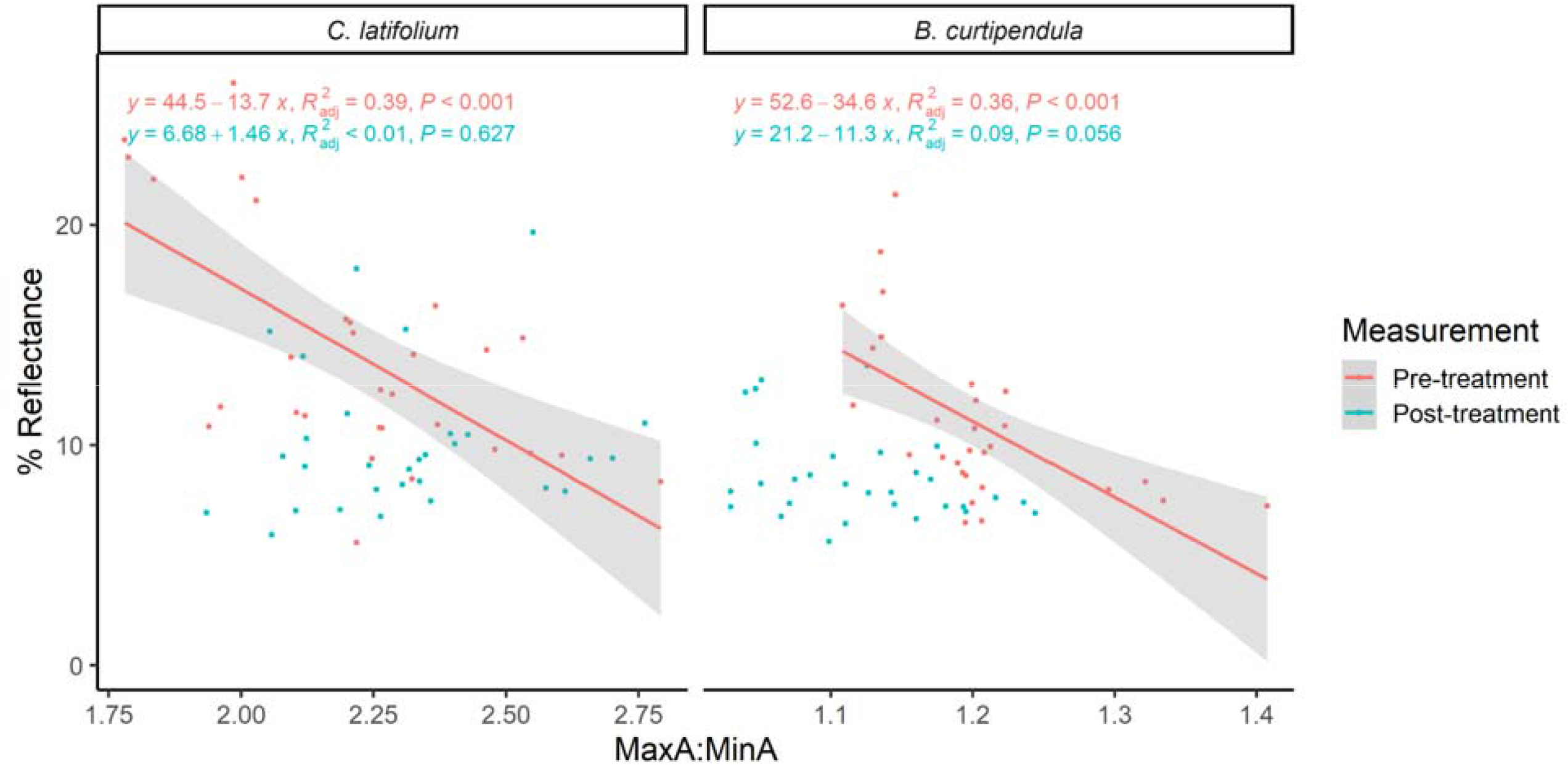

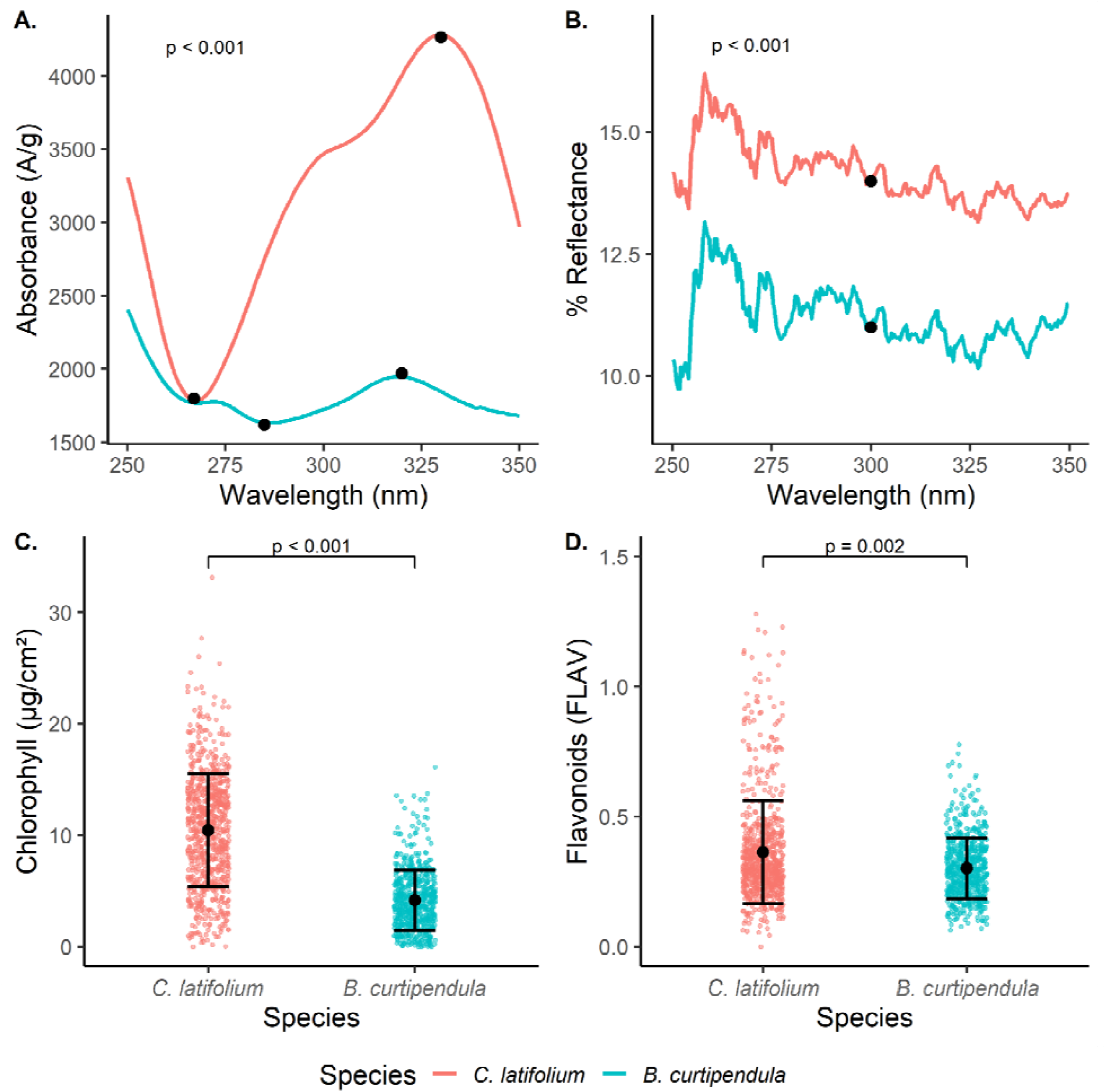

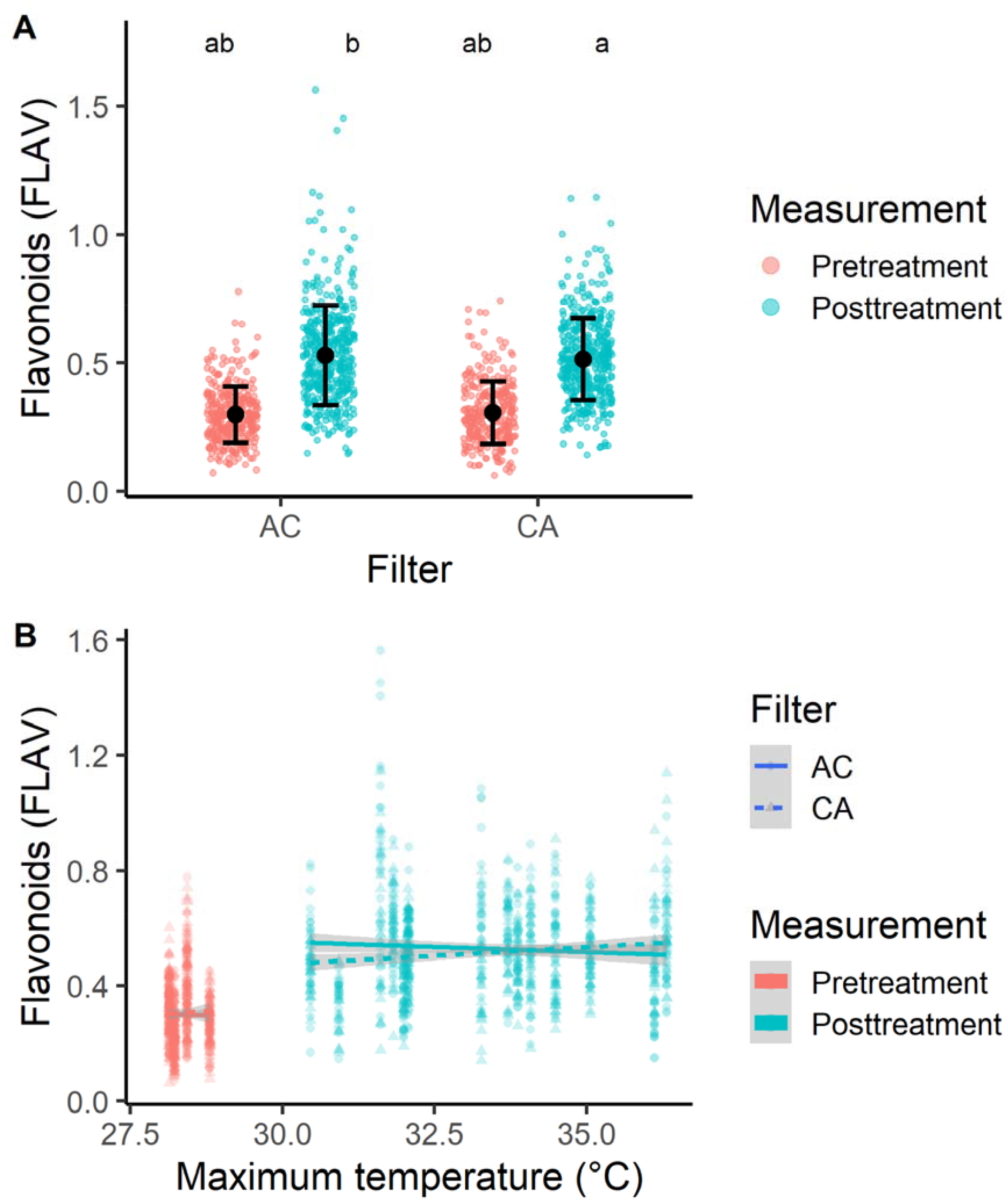

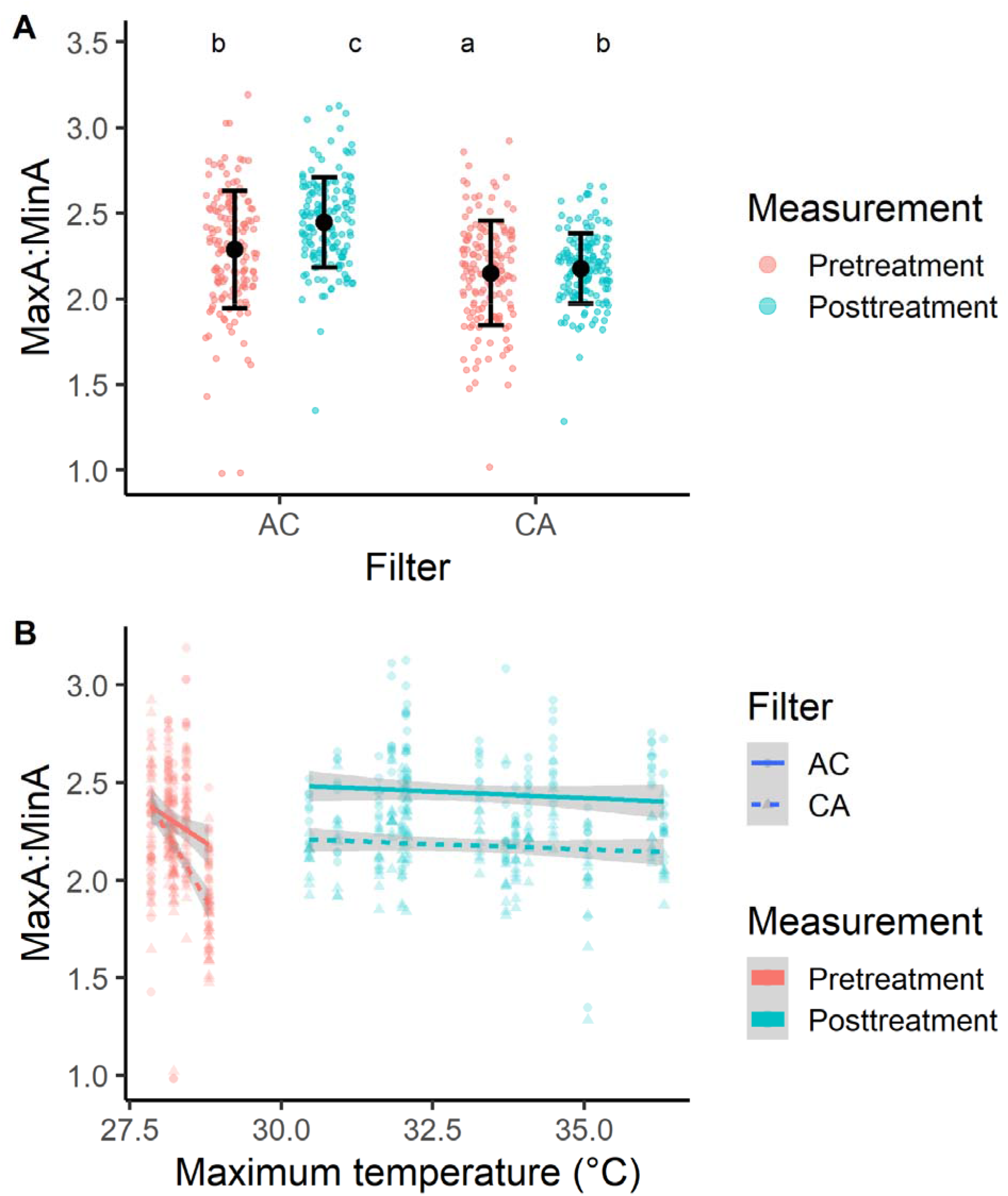

## Notes

### Competing Interest Statement

The authors have declared no competing interest.

## Works Cited

Balakumar, T., V. H. B. Vincent, and K. Paliwal. 1993. On the interaction of UV-B radiation (280–315 nm) with water stress in crop plants. Physiologia Plantarum 1:217–222. https://doi.org/10.1111/j.1399-3054.1993.tb00145.x

Ballaré, C. L., M. M. Caldwell, S.D. Flint, S. A. Robinson, J. F. Bornman. 2011. Effects of solar ultraviolet radiation on terrestrial ecosystems. Patterns, mechanisms, and interactions with climate change. Photochemical & Biological Sciences 1:225–241. https://doi.org/10.1039/C0PP90035D

Barnes, J. D., K. E. Percy, N. D. Paul, P. Jones, C. K. McLaughlin, P. M. Mullineaux, G. Creissen, and A. R. Wellburn. 1996a. The influence of UV-B radiation on the physicochemical nature of tobacco (Nicotiana tabacum L.) leaf surfaces. Journal of Experimental Botany 1:99–109. https://doi.org/10.1093/jxb/47.1.99

Barnes, P. W., A. R. Kersting, S. D. Flint, W. Beyschlag, and R. J. Ryel. 2013. Adjustments in epidermal UV-transmittance of leaves in sun-shade transitions. Physiologia Plantarum 1:200–213. https://doi.org/10.1111/ppl.12025

Barnes, P. W., C. L. Ballaré, and M. M. Caldwell. 1996b. Photomorphogenic Effects of UV-B Radiation on Plants: Consequences for Light Competition. Journal of Plant Physiology 1:15–20. https://doi.org/10.1016/S0176-1617(96)80288-4

Barnes, P. W., M. A. Tobler, K. Keefover-Ring, S. D. Flint, A. E. Barkley, R. J. Ryel, and R. L. Lindroth. 2016. Rapid modulation of ultraviolet shielding in plants is influenced by solar ultraviolet radiation and linked to alterations in flavonoids. Plant, Cell & Environment 1:222–230. https://doi.org/10.1111/pce.12609

Barnes, P. W., S. D. Flint, R. J. Ryel, M. A. Tobler, A. E. Barkley, and J. J. Wargent. 2015. Rediscovering leaf optical properties: New insights into plant acclimation to solar UV radiation. Plant Physiology and Biochemistry 1:94–100. https://doi.org/10.1016/j.plaphy.2014.11.015

Barr, D. J., R. Levy, C. Scheepers, and H. J. Tily. 2013. Random effects structure for confirmatory hypothesis testing: Keep it maximal. Journal of Memory and Language 1:255–278.

Basiouny, F. M., T. K. Van, and R. H. Biggs. 1978. Some Morphological and Biochemical Characteristics of C3 and C4 Plants Irradiated with UV-B. Physiologia Plantarum 1:29–32. https://doi.org/10.1111/j.1399-3054.1978.tb01533.x

Bidel, L. P. R., S. Meyer, Y. Goulas, Y. Cadot, and Z. G. Cerovic. 2007. Responses of epidermal phenolic compounds to light acclimation: In vivo qualitative and quantitative assessment using chlorophyll fluorescence excitation spectra in leaves of three woody species. Journal of Photochemistry and Photobiology B: Biology 1:163–179. https://doi.org/10.1016/j.jphotobiol.2007.06.002

Bilger, W., T. Johnsen, and U. Schreiber. 2001. UV-excited chlorophyll fluorescence as a tool for the assessment of UV-protection by the epidermis of plants. Journal of Experimental Botany 1:2007–2014. https://doi.org/10.1093/jexbot/52.363.2007

Blumthaler, M. 1993. Solar UV Measurements. Pages 71–94 in M., Tevini, editor. UV-B Radiation and Ozone Depletion: Effects on Humans, Animals, Plants, Microorganisms, and Materials, Lewis Publishers, Boca Raton, Florida.

Bokszczanin, K. L., S. Fragkostefanakis, H. Bostan, A. Bovy, P. Chaturvedi, M. L. Chiusano, N. Firon, R. Iannacone, S. Jegadeesan, K. Klaczynskid, H. Li, C. Mariani, F. Müller, P. Paul, M. Paupiere, E. Pressman, I. Rieu, K. D. Scharf, E. Schleiff, A. W. Van Heusden, W. Vriezen, W. Weckwerth, and P. Winter. 2013. Perspectives on deciphering mechanisms underlying plant heat stress response and thermotolerance. Frontiers in Plant Science 4. https://doi.org/10.3389/fpls.2013.00315

Caldwell, M. M., J. F. Bornman, C. L. Ballaré, S. D. Flint, and G. Kulandaivelu. 2007. terrestrial ecosystems, increased solar ultraviolet radiation, and interactions with other climate change factors. Photochemical & Photobiological Sciences 1:252–266. https://doi.org/10.1039/B700019G

Callaghan, T. V., L. O. Björn, Y. Chernov, T. Chapin, T. R. Christensen, B. Huntley, R. A. Ims, M. Johansson, D. Jolly, S. Jonasson, N. Matveyeva, N. Panikov, W. Oechel, G. Shaver, J. Elster, I. S. Jónsdóttir, K. Laine, K. Taulavuori, E. Taulavuori, and C. Zöckler. 2004. Responses to Projected Changes in Climate and UV-B at the Species Level. AMBIO: A Journal of the Human Environment 1:418–435. https://doi.org/10.1579/0044-7447-33.7.418

Casati, P., M. V. Lara, and C. S. Andreo. 2002. Regulation of enzymes involved in C4 photosynthesis and the antioxidant metabolism by UV-B radiation in Egeria densa, a submersed aquatic species. Photosynthesis Research 1:251. https://doi.org/10.1023/A:1015543208552

Cerovic, Z. G, G. Masdoumier, N. B. Ghozlen, and G. Latouche. 2012. https://onlinelibrary.wiley.com/doi/full/10.1111/j.1399-3054.2012.01639.x. Physiologia Plantarum 146(3):p251–260. https://doi.org/10.1111/j.1399-3054.2012.01639.x

Chipperfield, M. P., S. Bekki, S. Dhomse, N. R. P. Harris, B. Hassler, R. Hossaini, W. Steinbrecht, R. Thiéblemont, and M. Weber. 2017. Detecting recovery of the stratospheric ozone layer. Nature 549, 211–218. https://doi.org/10.1038/nature23681

Coffey, A., and M. A. K. Jansen. 2019. Effects of natural solar UV-B radiation on three Arabidopsis accessions are strongly affected by seasonal weather conditions. Plant Physiology and Biochemistry 1:64–72. https://doi.org/10.1016/j.plaphy.2018.06.016

Conner, J. K., and R. Neumeier. 2002. The effects of ultraviolet-B radiation and intraspecific competition on growth, pollination success, and lifetime female fitness in Phacelia campanularia and P. purshii (Hydrophyllaceae). American Journal of Botany 1:103–110. https://doi.org/10.3732/ajb.89.1.103

Correia, C. M., J. M. M. Pereira, J. F. Coutinho, L. O. Björn, and J. M. G. Torres-Pereira. 2005. Ultraviolet-B radiation and nitrogen affect the photosynthesis of maize: a Mediterranean field study. European Journal of Agronomy 1:337–347.

Csepregi, K., P. Teszlák, L. Kőrösi, and É. Hideg. 2019. Changes in grapevine leaf phenolic profiles during the day are temperature rather than irradiance driven. Plant Physiology and Biochemistry 1:169–178.

Flint, S., M. M. Caldwell. 2003. A biological spectral weighting function for ozone depletion research with higher plants. Physiologia Plantarum 1:137–144. https://doi.org/10.1034/j.1399-3054.2003.1170117.x

Flint, S. D., and M. M. Caldwell. 2002. Solar UV-B and visible radiation in tropical forest gaps: measurements partitioning direct and diffuse radiation. Global Change Biology 1:863–870. https://doi.org/10.1046/j.1365-2486.1998.00191.x

Gardner, G., C. Lin, E. M. Tobin, H. Loehrer, and D. Brinkman. 2009. Photobiological properties of the inhibition of etiolated Arabidopsis seedling growth by ultraviolet-B irradiation. Plant, Cell & Environment 1:1573–1583. https://doi.org/10.1111/j.1365-3040.2009.02021.x

Hyyryläinen, A., P. Rautio, M. Turunen, and S. Huttunen. 2015. Seasonal and inter-annual variation in the chlorophyll content of three co-existing Sphagnum species exceeds the effect of solar UV reduction in a subarctic peatland. SpringerPlus 1:478. https://doi.org/10.1186/s40064-015-1253-7

Izaguirre, M. M., C. A. Mazza, A. SvatoŠ, I. T. Baldwin, and C. L. BallarÉ. 2007. Solar Ultraviolet-B Radiation and Insect Herbivory Trigger Partially Overlapping Phenolic Responses in Nicotiana attenuata and Nicotiana longiflora. Annals of Botany 1:103–109.

Jansen, M. A. K., V. Gaba, and B. M. Greenberg. 1998. Higher plants and UV-B radiation: balancing damage, repair and acclimation. Trends in Plant Science 1:131–135. https://doi.org/10.1016/S1360-1385(98)01215-1

Jenkins, G. I. 2009. Signal Transduction in Responses to UV-B Radiation. Annual Review of Plant Biology 1:407–431. https://doi.org/10.1146/annurev.arplant.59.032607.092953

Kalbina, I., S. Li, G. Kalbin, L. O. Bjorn, A. Strid. 2008. Two separate UV-B radiation wavelength regions control expression of different molecular markers in Arabidopsis thaliana. Functional Plant Biology 1:222–227. https://doi.org/10.1071/FP07197

Kataria, S., K. N. Guruprasad, S. Ahuja, and B. Singh. 2013. Enhancement of growth, photosynthetic performance and yield by exclusion of ambient UV components in C3 and C4 plants. Journal of Photochemistry and Photobiology B: Biology 1:140–152. https://doi.org/10.1016/j.jphotobiol.2013.08.013

Kerr, J. B., C. T. McElrow. 1993. Evidence for large upward trends of ultraviolet-B radiation linked to ozone depletion. Science 262.5136:1032–1034. https://doi.org/10.1126/science.262.5136.1032

Kollias, N., A. H. Baqer, H. Ou-Yang. 2003. Diurnal and seasonal variations of the UV cut-off wavelength and most erythemally effective wavelength of solar spectra. Photodermatology, Photoimmunology & Photomedicine 1:89–92. https://doi.org/10.1034/j.1600-0781.2003.00002.x

Krizek, D. T. 2004. Influence of PAR and UV-A in Determining Plant Sensitivity and Photomorphogenic Responses to UV-B Radiation. Photochemistry and Photobiology 1:307–315. https://doi.org/10.1111/j.1751-1097.2004.tb00013.x

Lefcheck, J. S. 2016. piecewiseSEM: Piecewise structural equation modelling in r for ecology, evolution, and systematics. Methods in Ecology and Evolution 1:573–579.

Li, J., T. M. Ou-Lee, R. Raba, R. G. Amundson, and R. L. Last. 1993. Arabidopsis Flavonoid Mutants Are Hypersensitive to UV-B Irradiation. The Plant Cell 1:171–179. https://doi.org/10.1105/tpc.5.2.171

Li, X. X. Ke, H. Zhou, and Y. Tang. 2019. Contrasting altitudinal patterns of leaf UV reflectance and absorbance in four herbaceous species on the Qinghai–Tibetan Plateau. Journal of Plant Ecology, 1:245–254. https://doi.org/10.1093/jpe/rty016

Mancinelli, A. L., and A. Tolkowsky. 1968. Phytochrome and Seed Germination. V. Changes of Phytochrome Content During the Germination of Cucumber Seeds. Plant Physiology 1:489–494. https://doi.org/10.1104/pp.43.4.489

Martz, F., M. Turunen, R. Julkunen-Tiitto, H. Suokanerva, and M. Sutinen. 2011. Different response of two reindeer forage plants to enhanced UV-B radiation: modification of the phenolic composition. Polar Biology 1:411–420. https://doi.org/10.1007/s00300-010-0896-7

Mazza, C. A., H. E. Boccalandro, C. V. Giordano, D. Battista, A. L. Scopel, and C. L. Ballaré. 2000. Functional Significance and Induction by Solar Radiation of Ultraviolet-Absorbing Sunscreens in Field-Grown Soybean Crops. Plant Physiology 1:117–126. https://doi.org/10.1104/pp.122.1.117

McArthur, J. A., and W. R. Briggs. 1970. Phytochrome appearance and distribution in the embryonic axis and seedling of Alaska peas. Planta 1:146–154. https://doi.org/10.1007/BF00386098

McDonald, G. K., and G. M. Paulsen. 1997. High temperature effects on photosynthesis and water relations of grain legumes. Plant and Soil 1:47–58.

Mori, K., N. Goto-Yamamoto, M. Kitayama, and K. Hashizume. 2007. Loss of anthocyanins in red-wine grape under high temperature. Journal of Experimental Botany 1:1935–1945.

Mori, K., S. Sugaya, and H. Gemma. 2005. Decreased anthocyanin biosynthesis in grape berries grown under elevated night temperature condition. Scientia Horticulturae 1:319–330.

Muggeo, M. R. 2003. Estimating regression models with unknown break-points. Statistics in Medicine 22(19):3055–3071. https://doi.org/10.1002/sim.1545

Pal, M., P. H. Zaidi, S. R. Voleti, and A. Raj. 2006. Solar UV-B Exclusion Effects on Growth and Photosynthetic Characteristics of Wheat and Pea. Journal of New Seeds 1:19–34. https://doi.org/10.1300/J153v08n01_02

Robson, T. M., P. J. Aphalo, A. Katarzyna Banaś, P. W. Barnes, C. C. Brelsford, G. I. Jenkins, T. K. Kotilainen, J. Łabuz, J. Martínez-Abaigar, L. O. Morales, S. Neugart, M. Pieristè, N. Rai, F. Vandenbussche, and M. A. K. Jansen. 2019. A perspective on ecologically relevant plant-UV research and its practical application. Photochemical & Photobiological Sciences 1:970–988. https://doi.org/10.1039/C8PP00526E

Robson, T. M., V. A. Pancotto, S. D. Flint, C. L. Ballaré, O. E. Sala, A. L. Scopel, and M. M. Caldwell. 2003. Six years of solar UV-B manipulations affect growth of Sphagnum and vascular plants in a Tierra del Fuego peatland. New Phytologist 1:379–389. https://doi.org/10.1046/j.1469-8137.2003.00898.x

Ros, J. and M. Tevini. Interaction of UV-Radiation and IAA During Growth of Seedlings and Hypocotyl Segments of Sunflower. 1995. Journal of Plant Physiology: 1:295–302. https://doi.org/10.1016/S0176-1617(11)82057-2

Sampson, B. J., and J. H. Cane. 1999. Impact of enhanced ultraviolet-B radiation on flower, pollen, and nectar production. American Journal of Botany 1:108–114. https://doi.org/10.2307/2656959

Sayed, O. H., M. J. Earnshaw, and M. J. Emes. 1989. Photosynthetic Responses of Different Varieties of Wheat to High Temperature: II. Effect of Heat Stress on Photosynthetic Electron Transport. Journal of Experimental Botany 1:633–638.

Searles, P. S., B. R. Kropp, S. D. Flint, and M. M. Caldwell. 2001. Influence of solar UV-B radiation on peatland microbial communities of southern Argentinia. New Phytologist 1:213–221. https://doi.org/10.1046/j.0028-646X.2001.00254.x

Shepherd, T., and D. W. Griffiths. 2006. The effects of stress on plant cuticular waxes. New Phytologist 1:469–499. https://doi.org/10.1111/j.1469-8137.2006.01826.x

Shinkle, J. R., A. K. Atkins, E. E. Humphrey, C. W. Rodgers, S. L. Wheeler, and P. W. Barnes. 2004. Growth and morphological responses to different UV wavebands in cucumber (Cucumis sativum) and other dicotyledonous seedlings. Physiologia Plantarum 1:240–248. https://doi.org/10.1111/j.0031-9317.2004.0237.x

Shinkle, J. R., D. L. Derickson, P. W. Barnes. 2005. Comparative Photobiology of Growth Responses to Two UV-B Wavebands and UV-C in Dimred-light- and White-light-grown Cucumber (Cucumis sativus) Seedlings: Physiological Evidence for Photoreactivation. Photochemistry and Photobiology 1:1069–1074. https://doi.org/10.1562/2005-01-10-RA-411

Shinkle, J. R., M. C. Edwards, A. Koenig, A. Shaltz, and P. W. Barnes. 2010. Photomorphogenic regulation of increases in UV-absorbing pigments in cucumber (Cucumis sativus) and Arabidopsis thaliana seedlings induced by different UV-B and UV-C wavebands. Physiologia Plantarum 1:113–121. https://doi.org/10.1111/j.1399-3054.2009.01298.x

Smith, J. L., D. J. Burritt, and P. Bannister. 2000. Shoot Dry Weight, Chlorophyll and UV-B-absorbing Compounds as Indicators of a Plant’s Sensitivity to UV-B Radiation. Annals of Botany 1:1057–1063. https://doi.org/10.1006/anbo.2000.1270

Strømme, C. B., R. Julkunen-Tiitto, U. Krishna, A. Lavola, J. E. Olsen, and L. Nybakken. 2015. UV-B and temperature enhancement affect spring and autumn phenology in Populus tremula. Plant, Cell & Environment 1:867–877. https://doi.org/10.1111/pce.12338

teramura, A. H., and J. H. Sullivan. 1994. Effects of UV-B radiation on photosynthesis and growth of terrestrial plants. Photosynthesis Research 1:463–473. https://doi.org/10.1007/BF00014599

tilbrook, K., A. B. Arongaus, M. Binkert, M. Heijde, R. Yin, and R. Ulm. 2013. The UVR8 UV-B Photoreceptor: Perception, Signaling and Response. The Arabidopsis Book / American Society of Plant Biologists 11. https://doi.org/10.1199/tab.0164

tobin, E. M., and W. R. Briggs. 1969. Phytochrome in Embryos of Pinus palustris1. Plant Physiology 1:148–150. https://doi.org/10.1104/pp.44.1.148

tossi, V. E., J. J. Regalado, J. Iannicelli, L. E. Laino, H. P. Burrieza, A. S. Escandón, and S. I. Pitta-Álvarez. 2019. Beyond Arabidopsis: Differential UV-B Response Mediated by UVR8 in Diverse Species. Frontiers in Plant Science 10. https://doi.org/10.3389/fpls.2019.00780

Ulm, R., A. Baumann, A. Oravecz, Z. Máté, É. Ádám, E. J. Oakeley, E. Schäfer, and F. Nagy. 2004. Genome-wide analysis of gene expression reveals function of the bZIP transcription factor HY5 in the UV-B response of Arabidopsis. Proceedings of the National Academy of Sciences 1:1397–1402. https://doi.org/10.1073/pnas.0308044100

Vito M. R. Muggeo (2008). segmented: an R Package to Fit Regression Models with Broken-Line Relationships. R News, 8/1, 20-25. URL https://cran.r-project.org/doc/Rnews/.

Yao, X., and Q. Liu. 2006. Changes in morphological, photosynthetic and physiological responses of Mono Maple seedlings to enhanced UV-B and to nitrogen addition. Plant Growth Regulation 1:165. https://doi.org/10.1007/s10725-006-9116-4

