## Supplementary material for "Natural Shortwave UV-B Radiation Increases UV absorbing pigments levels in Texas native grasses *Chasmanthium latifolium* and *Bouteloua curtipendula*": All supplements

Supplemental Materials

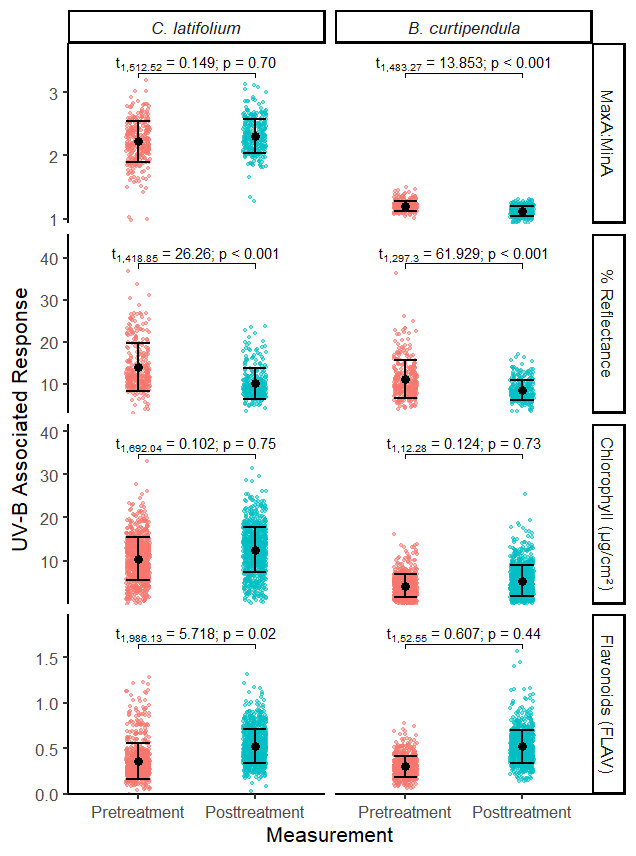

*Figure S1*. Impact of exposure to the field environment on *C. latifolium* and *B. curtipendula* UV-B associated responses (pretreatment in red, posttreatment in blue). Shown is a Post-hoc Tukey's honest significance test of the estimated marginal means, datapoints, and mean ± SD.

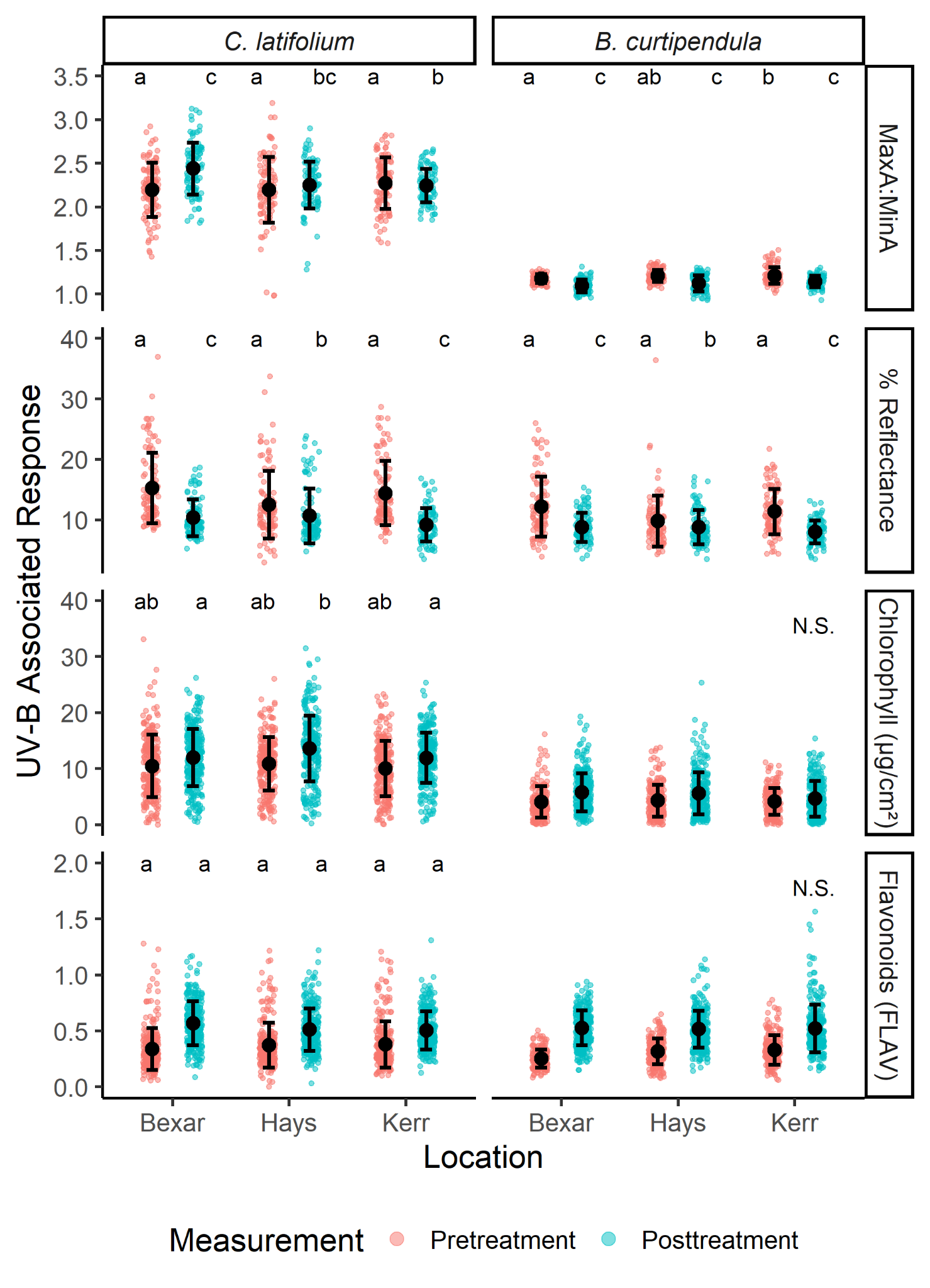

*Figure S2*. Impact of exposure to the field environment on *C. latifolium* and *B. curtipendula* UV-B associated responses by location (Bexar, Hays, Kerr) and measurement (pretreatment in red, posttreatment in blue). Shown are significance groups (treatments with the same letter code are not significantly different; Post-hoc Tukey's honest significance test difference test of estimated marginal means, p > 0.05), datapoints, and mean ± SD.

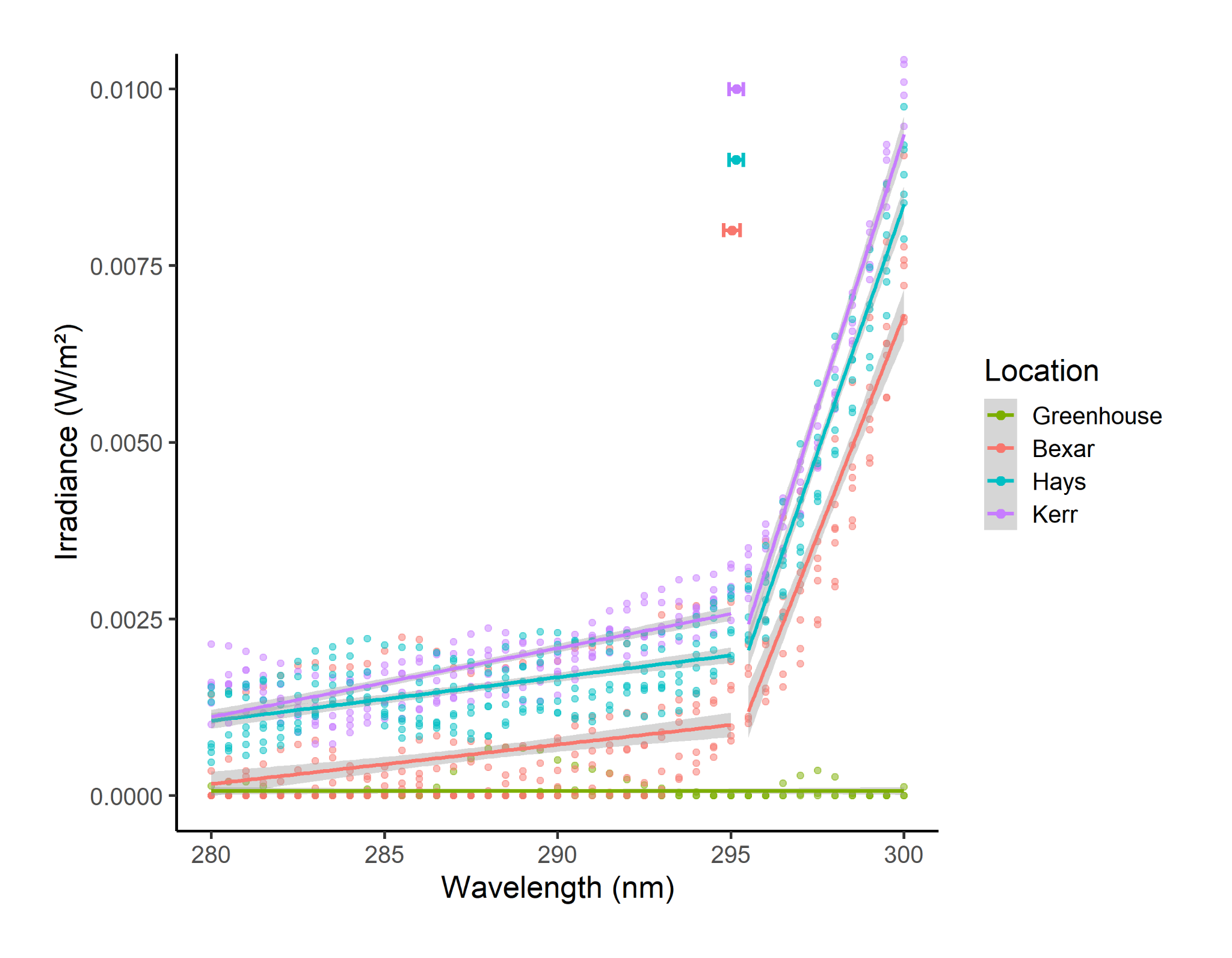

*Figure S3*. Solar irradiance spectra in W/m^2^ across the SW-UVB spectrum (280-300 nm) at Bexar County (red), Kerr County (purple), Hays County (blue), and the Trinity University greenhouse (green) at solar noon. Shown are raw data points, lines of best fit, and 95% confidence intervals of the lines of best fit (grey bars) and breakpoints (colored dots and bars). We conducted a breakpoint analysis with R package segmented (Vito and Muggeo 2008), by first utilizing a pseudo Score test to detect the presence of a breakpoint at each site, then located the breakpoint location through iterative linear models as described in Muggeo (2003). The final segmented linear model accounted for (adjusted R^2^) 95.8% of the variation in irradiance values between 280 and 300 nm, with the Bexar (295.030 nm (294.791–295.269 nm); p < 2×10^-16^), Hays (295.150 nm (294.944–295.357 nm); p < 2×10^-16^), and Kerr fieldsites (295.156 nm (294.953–295.359 nm); p < 2×10^-16^) containing a singular breakpoint, while the greenhouse exhibited consistently low irradiance throughout the SW-UVB spectra (298.373 nm (293.188–303.558 nm); p = 0.45).

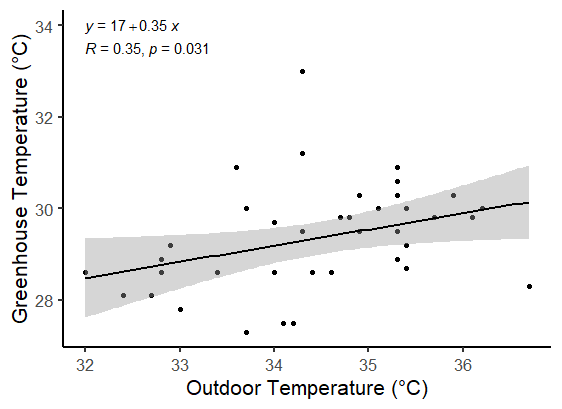

*Figure S4*. Relationship between outdoor and greenhouse temperature (°C; n = 39). Shown are the line of best fit equation and Pearson’s correlation values, data, line of best fit, and 95% confidence interval.

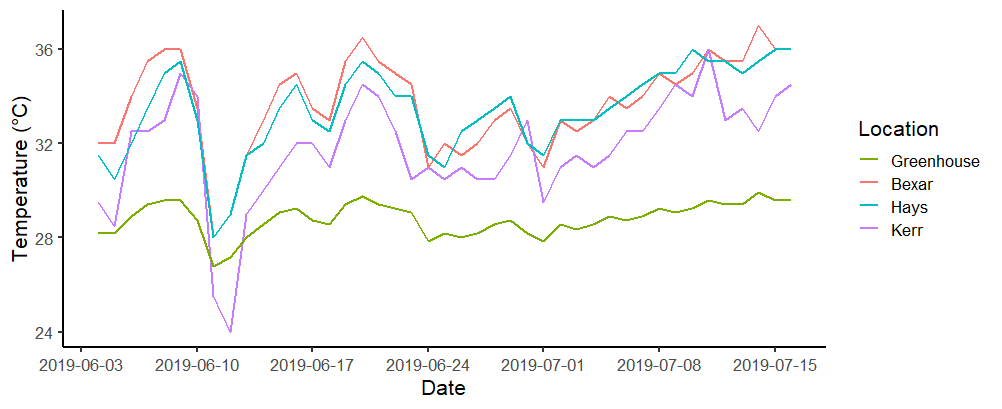

*Figure S5*. Maximum daily temperature (°C) recorded at Bexar, Hays, and Kerr sites and estimated maximum daily greenhouse temperature (°C) from June 4, 2019 to July 16, 2019.

*Table S1*. Post-hoc Tukey's honest significance test difference test results comparing the estimated marginal means of *C. latifolium and B. curtipendula* UV-B associated responses by location (Bexar, Hays, and Kerr) and measurement (pretreatment and posttreatment).

| Comparison | Result | UV-B Associated Response | | | | | |
| --- | --- | --- | --- | --- | --- | --- | --- |
|  |  | *C. latifolium* | | | | *B. curtipendula* | |
|  |  | MaxA:MinA | % Reflectance | Chlorophyll (µg/cm^2^) | Flavonoids (FLAV) | MaxA:MinA | % Reflectance |
| Bexar Posttreatment - Hays Posttreatment | df | 28.9 | 51.4 | 46.6 | 31.1 | 35.3 | 43 |
|  | t | 3.004 | -0.329 | -3.341 | 2.317 | -2.865 | 0.026 |
|  | p | 0.06 | 1.00 | 0.02 | 0.22 | 0.07 | 1.00 |
| Bexar Posttreatment - Kerr Posttreatment | df | 33.6 | 73.3 | 67.5 | 38.9 | 45.1 | 57 |
|  | t | 4.516 | -1.484 | 0.23 | 1.608 | -2.307 | 0.419 |
|  | p | 0.001 | 0.68 | 1.00 | 0.60 | 0.21 | 1.00 |
| Bexar Pretreatment - Bexar Posttreatment | df | 506.6 | 565.9 | 1457.8 | 1390.7 | 552.9 | 334.6 |
|  | t | -7.943 | 13.231 | -1.663 | 0.86 | -6.518 | 14.188 |
|  | p | < 0.001 | < 0.001 | 0.56 | 0.96 | < 0.001 | < 0.001 |
| Bexar Pretreatment - Hays Posttreatment | df | 431.2 | 587.2 | 1439.7 | 1156.4 | 548.9 | 350.8 |
|  | t | -6.333 | 13.042 | -2.379 | 1.514 | -7.243 | 13.986 |
|  | p | < 0.001 | < 0.001 | 0.16 | 0.66 | < 0.001 | < 0.001 |
| Bexar Pretreatment - Hays Pretreatment | df | 28.9 | 51.3 | 56.6 | 35.2 | 35.3 | 42.9 |
|  | t | 0.004 | 4.941 | -0.832 | -1.272 | -2.949 | 5.066 |
|  | p | 1.00 | < 0.001 | 0.96 | 0.80 | 0.06 | < 0.001 |
| Bexar Pretreatment - Kerr Posttreatment | df | 423.4 | 586.5 | 1396.7 | 1127.2 | 545.4 | 310.7 |
|  | t | -5.612 | 12.557 | -1.572 | 1.313 | -7.031 | 13.95 |
|  | p | < 0.001 | < 0.001 | 0.62 | 0.78 | < 0.001 | < 0.001 |
| Bexar Pretreatment - Kerr Pretreatment | df | 28.9 | 51.3 | 56.8 | 34.9 | 35.3 | 42.9 |
|  | t | -1.236 | 1.51 | 0.763 | -1.574 | -3.29 | 1.688 |
|  | p | 0.82 | 0.66 | 0.97 | 0.62 | 0.03 | 0.55 |
| Hays Posttreatment - Kerr Posttreatment | df | 34.3 | 76.8 | 72.2 | 39.8 | 46.6 | 59.2 |
|  | t | 1.62 | -1.17 | 3.195 | -0.573 | 0.377 | 0.391 |
|  | p | 0.59 | 0.85 | 0.02 | 0.99 | 1.00 | 1.00 |
| Hays Pretreatment - Bexar Posttreatment | df | 430.6 | 587.1 | 1437.2 | 1155.4 | 548.6 | 348.1 |
|  | t | -7.5 | 11.878 | -1.463 | 1.22 | -5.413 | 12.578 |
|  | p | < 0.001 | < 0.001 | 0.69 | 0.83 | < 0.001 | < 0.001 |
| Hays Pretreatment - Hays Posttreatment | df | 507.8 | 565.9 | 1458.9 | 1395 | 553.2 | 337.3 |
|  | t | -6.71 | 11.9 | -2.206 | 1.949 | -6.494 | 12.781 |
|  | p | < 0.001 | < 0.001 | 0.24 | 0.37 | < 0.001 | < 0.001 |
| Hays Pretreatment - Kerr Posttreatment | df | 423.4 | 586.5 | 1396.3 | 1128 | 545.4 | 310.7 |
|  | t | -5.613 | 11.331 | -1.385 | 1.69 | -6.115 | 12.568 |
|  | p | < 0.001 | < 0.001 | 0.74 | 0.54 | < 0.001 | < 0.001 |
| Hays Pretreatment - Kerr Pretreatment | df | 28.9 | 51.3 | 54.9 | 35.1 | 35.3 | 42.9 |
|  | t | -1.24 | -3.431 | 1.607 | -0.299 | -0.34 | -3.378 |
|  | p | 0.81 | 0.01 | 0.60 | 1.00 | 1.00 | 0.02 |
| Kerr Pretreatment - Bexar Posttreatment | df | 430.6 | 587.1 | 1435.7 | 1153.9 | 548.6 | 348.1 |
|  | t | -7.018 | 12.74 | -1.832 | 1.311 | -5.306 | 13.509 |
|  | p | < 0.001 | < 0.001 | 0.45 | 0.78 | < 0.001 | < 0.001 |
| Kerr Pretreatment - Hays Posttreatment | df | 431.2 | 587.2 | 1438 | 1155.6 | 548.9 | 350.8 |
|  | t | -5.852 | 12.662 | -2.561 | 1.989 | -6.209 | 13.52 |
|  | p | < 0.001 | < 0.001 | 0.11 | 0.35 | < 0.001 | < 0.001 |
| Kerr Pretreatment - Kerr Posttreatment | df | 488.2 | 565.7 | 1412.4 | 1326.8 | 547.5 | 297.9 |
|  | t | -5.434 | 12.283 | -1.761 | 1.827 | -6.174 | 13.691 |
|  | p | < 0.001 | < 0.001 | 0.49 | 0.45 | < 0.001 | < 0.001 |
